# Towards understanding interindividual differences in cortical morphological brain networks

**DOI:** 10.1101/2020.12.21.423884

**Authors:** Zhen Li, Junle Li, Ningkai Wang, Jinhui Wang

## Abstract

Individual-level morphological brain networks are becoming an important approach for studying human connectome; however, their interindividual differences are not well understood with respect to behavioral and cognitive relevance, individual identification, and genetic origin. Using three publicly available datasets that involved cross-sectional and longitudinal structural magnetic resonance scans of adults and children, we constructed four morphological brain networks for each of 1,451 images from 1,329 participants on the basis of cerebral surface-based, vertex-wise cortical thickness, fractal dimension, gyrification index and sulcal depth, respectively. The morphological index-dependent networks were further fused via multiplex network model, and fed into community detection. We found that the multiplex morphological brain networks 1) accounted for significant proportions of interindividual variance in and were predictive of multiple behavioral and cognitive domains, in particular Cognition and Motor domains (*P* < 0.05, corrected), 2) distinguished individuals from each other with high accuracies even for twin subjects (accuracies > 96%), and 3) exhibited low-moderate heritability with the highest for sulcal depth-based morphological brain networks. Intriguingly, compared with intra-module morphological connectivity, inter-module connections explained more behavioral and cognitive variance and were associated with higher heritability. Further comparisons revealed that multiplex morphological brain networks outperformed each type of single-layer morphological brain networks in the performance of behavioral and cognitive association and prediction, and individual identification. Finally, all the findings were generally reproducible over different datasets. Altogether, our findings indicate that interindividual differences in individual-level morphological brain networks are biologically meaningful, which underpins their usage as fingerprints for individualized studies in health and disease.

## Introduction

Morphological covariance networks, which encode interregional covariance patterns in local brain morphology (e.g., cortical thickness) across a cohort of participants, have long been used to study cerebral architecture in health and disease (Alexander-Bloch, Giedd et al. 2013, Evans 2013). However, these population-based approaches lead to obscure neurobiological significance of the constructed morphological brain networks due to the neglect of interindividual differences. Recent methodological advances in constructing individual-level morphological similarity networks make it possible for neurobiologically understanding morphological brain networks by studying behavioral and cognitive relevance, and genetic origin of their interindividual differences.

Currently, there are several different approaches developed to construct morphological brain networks at the individual level (Tijms, Seriès et al. 2012, Kong, Liu et al. 2015, Wang, Jin et al. 2016, Li, Yang et al. 2017, Seidlitz, Vasa et al. 2018, Li et al., 2020). With these approaches, some pioneer studies have found that morphological brain networks are associated with specific behavioral and cognitive performance among individuals (Tijms, Yeung et al. 2014, Seidlitz, Vasa et al. 2018), correlated with individual ages (Kong, Liu et al. 2015, Corps and Rekik 2019), capable of distinguishing patients from controls (Mahjoub, Mahjoub et al. 2018, Zhao, Guo et al. 2020), and able to predict clinical progression of diseases (Tijms, Ten Kate et al. 2018). Despite these efforts, there are still several fundamental issues that remain elusive for interindividual differences in morphological brain networks, such as what do they originate from, and to what extent they determine various downstream phenotypes.

The advent of publicly available large-scale datasets (e.g., the Human Connectome Project and UK Biobank) that cover multidimensional data (e.g., brain imaging data, genetic information, and behavior and cognition) provides us opportunities to understand interindividual differences in brain structure and function. With these resources, previous studies have shown that structural and functional brain networks are under genetic control (e.g., Elliott, Sharp et al. 2018, Zhong, Wei et al. 2020), are related to and can predict a wide range of behavior and cognition (e.g., (Smith, Nichols et al. 2015, Liegeois, Li et al. 2019, Sripada, Angstadt et al. 2019), and can serve as fingerprints to identify individuals (e.g., (Finn, Shen et al. 2015, Miranda-Dominguez, Feczko et al. 2018). For morphological brain networks, however, there are no such studies that examine their genetic origin and whether they can act as alternative fingerprints to distinguish individuals from each other. Although there is preliminary evidence for behavioral and cognitive relevance of morphological brain networks (Tijms, Yeung et al. 2014, Seidlitz, Vasa et al. 2018), existing studies are mainly based on univariate analysis of a few discrete behavioral and cognitive variables using small sample size. There still lacks a holistic, multivariate examination of the extent to which morphological brain networks can account for and predict interindividual differences in various behavioral and cognitive domains from a large cohort of participants.

In this study, we aimed to provide convincing, cross-dataset validated evidence for neurobiological significance of morphological brain networks by examining their interindividual differences in the context of behavioral and cognitive relevance, individual identification, and genetic origin. To this end, we utilized three large-scale, publicly available datasets to construct individual multiplex morphological brain networks that integrated complementary information from different morphological indices. After identifying communities or modules embedded in the multiplex morphological brain networks, intra- and inter-module morphological connectivity were extracted and used to account for interindividual variance in multiple behavior and cognition domains with a multivariate variance component model (Ge, Reuter et al. 2016, Liegeois, Li et al. 2019), predict individual behavioral and cognitive performance with a multivariate brain basis set modeling method (Sripada, Angstadt et al. 2019), and identify individuals using a network matching method (Finn, Shen et al. 2015). Finally, an ACE model (Chen, Formisano et al. 2019) was used to examine the extent to which morphological brain networks were under genetic control. We hypothesized that morphological brain networks 1) can account for interindividual variance in behavior and cognition; 2) can predict individual behavior and cognition; 3) can identify individuals with high accuracy; and 4) are heritable.

## Materials and Methods

### General description of datasets and analytical pipeline

This study included three publicly available datasets: the Human Connectome Project (HCP) S1200 dataset (www.humanconnectome.org) (Van Essen, Smith et al. 2013), the Beijing Normal University (BNU) test-retest dataset (http://fcon_1000.projects.nitrc.org/indi/CoRR/html/bnu_1.html) (Lin, Dai et al. 2015), and the “Longitudinal Brain Correlates of Multisensory Lexical Processing in Children” (LBCMLPC) dataset (https://openneuro.org) (Lytle, McNorgan et al. 2019). For each T1-weight structural image from each participant in the datasets, we first derived four cerebral surface-based, vertex-wise morphological maps of cortical thickness (CT), fractal dimension (FD), gyrification index (GI), and sulcal depth (SD). The morphological maps were then used to construct morphological index-dependent, single-layer networks, which were further merged into multiplex morphological brain networks. After identifying community structure of the resultant single-layer and multiplex morphological brain networks at the group level, the HCP dataset was used to analyze behavioral and cognitive association and prediction (unrelated participants) and heritability (twin participants) of morphological brain networks, the BNU dataset was used to analyze the potential of morphological brain networks as fingerprints for individual identification, and the LBCMLPC dataset was used for validation of reproducibility (behavioral and cognitive association and prediction, and individual identification). Figure 1 presents a flowchart of overall data analytical pipeline of this study.

**Figure 1.**
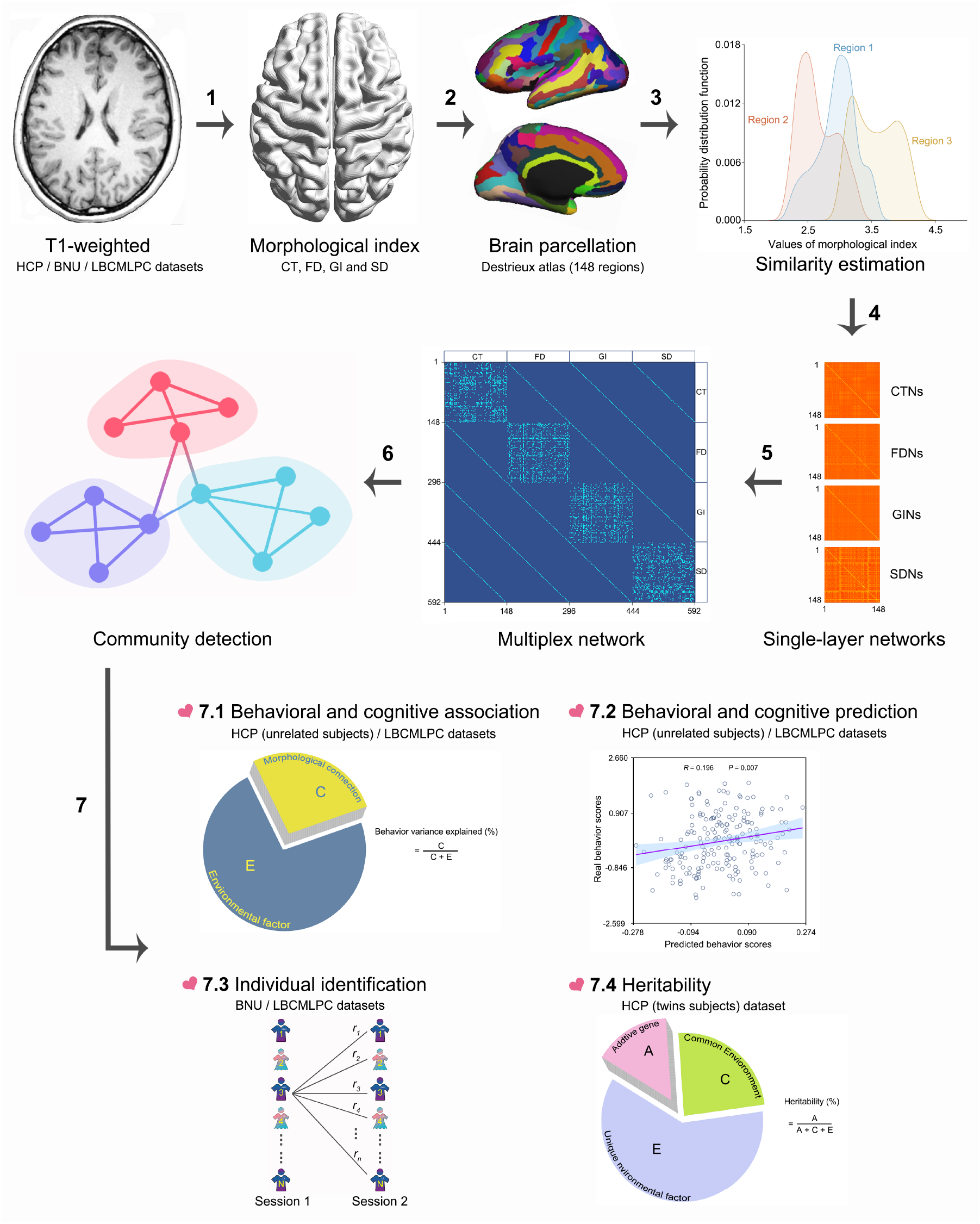
A flowchart of data analytical pipeline in this study. **1**) Each T1-weighted structural image in the HCP, BNU, and LBCMLPC datasets was processed to extract four cortical surface-based, vertex-wise morphological maps (CT, FD, GI and SD). **2**) The Destrieux atlas was used to divide the cortical surface into148 regions of interest. **3**) For each region, a probability distribution function was estimated based on the intraregional values of each morphological index. **4**) Interregional morphological connectivity was estimated between any pair of regions using the JSD-based similarity, resulting in four morphological index-dependent, single-layer morphological brain networks. **5**) The morphological index-dependent, single-layer networks were further fused to form a multiplex morphological brain network. **6**) Community detection was performed for the resultant single-layer and multiplex morphological brain networks at the group level. **7**) Based on the identified community structure, morphological connectivity within each module and between each pair of modules were extracted and used to examine their abilities to **7**.**1**) explain interindividual variance in behavior and cognition, **7**.**2**) predict individual performance of behavior and cognition, and **7**.**3**) distinguish individuals from each other, and **7**.**4**) their heritability. HCP, Human Connectome Project; BNU, Beijing Normal University; LBCMLPC, Longitudinal Brain Correlates of Multisensory Lexical Processing in Children; CT, cortical thickness; FD, fractal dimension; GI, gyrification index; SD, sulcal depth; CTNs, cortical thickness networks; FDNs, fractal dimension networks; GINs, gyrification index networks; SDNs, sulcal depth networks.

### Participants and data acquisition

#### HCP dataset

Out of 1113 subjects with T1-weight structural images in the HCP dataset, a total of 650 unrelated healthy participants and 217 pairs of monozygotic (MZ) and dizygotic (DZ) twins (age, 22-35 years) were included in this study. T1-weighted structural images were acquired from each participant on a Siemens Skyra Connectome scanner with a magnetization prepared rapid gradient echo (MPRAGE) sequence: repetition time (TR) = 2400 ms; inversion time (TI) = 1000 ms; echo time (TE) = 2.14 ms; voxel size = 0.7 × 0.7 × 0.7 mm^3^; field of view (FOV) = 224 × 224 mm^2^; flip angle = 8°; bandwidth (BW) = 210 Hz/Px. Out of the 434 twin participants, 8 pairs completed two structural image scans.

#### BNU dataset

The BNU dataset contained 57 participants (age, 19-30 years) with no history of neurological and psychiatric disorders. Individual T1-weighted structural images were acquired on a 3T Siemens Trio-Tim MRI scanner using the MPRAGE sequence: TR = 2530 ms; TE = 3.39 ms; TI = 1100 ms; slice thickness = 1.33 mm; number of slices = 144; no interslice gap; FOV = 256 × 256 mm^2^; flip angle = 7°. All participants completed two scan sessions within an interval of approximate 6 weeks (40.94 ± 4.51 days).

#### LBCMLPC dataset

There were 188 children in the LBCMLPC dataset with no history of psychiatric illness or neurological disease. All participants were approximately 10.5 years old at data scan 1 of whom 49 participants returned approximately 2.5 years later for the second data scan. For each child, T1-weighted structural images were collected on a 3T Siemens Trio-Tim scanner using the MPRAGE sequence: TR = 2300 ms; TE = 3.36 ms; FOV = 256 × 256 mm^2^; slice thickness = 1 mm; number of slices = 160; voxel size = 1 × 1 × 1 mm^3^; flip angle = 9°; BW = 240 Hz/Px.

### Behavioral and cognitive measures

#### HCP dataset

The HCP dataset included a broad range of behavioral and cognitive measures that were mainly evaluated via the NIH Toolbox Assessment of Neurological and Behavioral function. Out of original 581 items, a total of 60 were screened out and included in this study (Supplemental Materials). These items belonged to six domains, including Alertness, Cognition, Emotion, Motor, Personality, and Sensory (Fig. 4A and Table S1). After standardizing each item to make the scale comparable (Z-score transformation across participants), the items were averaged within each domain and used for subsequent analyses.

#### LBCMLPC dataset

Each participant in the LBCMLPC dataset completed a series of standardized psycho-educational tests at data scan 1, including the Woodcock-Johnson III Test, Wechsler Abbreviated Scale for Intelligence, Comprehensive Test for Phonological Processing, and Test for Word Reading Efficiency. Out of original 50 items, a total of 26 were screened out and included in this study (Supplemental Materials; Fig. S6A and Table S2). Similar to the HCP dataset, the items were standardized across participants, and averaged within each domain.

### Structural image preprocessing

All structural images in the three datasets underwent the same standard preprocessing pipeline using the CAT12 toolbox (http://dbm.neuro.uni-jena.de/cat12/) based on the SPM12 package (https://www.fil.ion.ucl.ac.uk/spm/software/spm12/). The CAT12 toolbox offers a fast and reliable alternative to FreeSurfer for analysis of cerebral surface-based morphometry, and allows computing multiple indices, including CT, FD, GI and SD. Briefly, each structural image was first segmented into gray matter, white matter and cerebrospinal fluid based on an adaptive Maximum A Posterior technique (Rajapakse 1997). Then, CT was estimated using a projection-based thickness method (Dahnke, Yotter et al. 2013), and a central surface was created. For the resultant central surface, a topology correction based on spherical harmonics was used to account for topological defects (Yotter, Dahnke et al. 2011). Based on the spherical harmonic reconstructions, FD, GI and SD were calculated as the slope of a logarithmic plot of surface area versus the maximum *l*-value (Yotter, Nenadic et al. 2011), absolute mean curvature (Luders, Thompson et al. 2006), and Euclidean distance between the central surface and its convex hull, respectively. All computations of CT, FD, GI and SD were conducted in subject native space with default parameter settings. Finally, individual morphological maps of CT, FD, GI and SD were resampled into the common fsaverage template and smoothed using a Gaussian kernel (15-mm full width at half maximum for CT and 25-mm full width at half maximum for the others).

### Construction of single-layer morphological brain networks

In this study, we constructed four morphological index-dependent brain networks for each structural image by estimating interregional morphological similarity in terms of distributions of intraregional CT, FD, GI, and SD, respectively.

#### Brain parcellation

To definite brain regions, we utilized the Destrieux atlas (Destrieux, Fischl et al. 2010) to parcellate the cortex surface into 148 regions of interest, with each representing a node in the morphological brain networks.

#### Estimate of morphological similarity

To quantify interregional morphological similarity, we used a Jensen-Shannon divergence (JSD)-based approach as in our previous study (Li et al., 2020). Specifically, for each morphological index derived from each structural image, values of all vertices within each region were first extracted, based on which a probability density function was estimated using a kernel density function (MATLAB function, ksdensity). After converting the resultant probability density functions to probability distribution functions (PDFs), the JSD was then calculated between any pair of regions using their PDFs. Formally, for two PDFs *P* and *Q*, the JSD was calculated as:

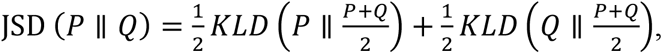

where 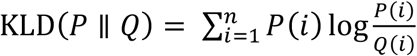 with *n* being the number of sample points (2^8^ in this study) (Wang, Jin et al. 2016). The JSD was further transformed to a distance metric (square root) and subtracted from 1, resulting in values that denoted the degree of similarity between two PDFs (0, absolutely different; 1, exactly the same). Finally, we obtained four 148 × 148 morphological brain networks for each structural image.

### Construction of multiplex morphological brain networks

For the morphological index-dependent brain networks derived above, our previous study showed that they exhibited low spatial similarities in overall connectivity patterns (Li et al., 2020). These findings were further replicated by this study in all datasets (see Results). Thus, to provide a more holistic understanding of the morphological brain networks, we employed a multiplex network model to merge the four morphological index-dependent brain networks derived from each structural image to a multiplex network (Betzel and Bassett 2017, De Domenico 2017). In the resultant multiple networks, the morphological index-dependent morphological brain networks were treated as different layers, which were interconnected via edges that linked each node in each layer with replicas of the node in the other layers. For the inter-layer edges, their weights were defined as JSD-based similarities in regional PDFs derived from different morphological indices of the same regions.

### Community detection

We detected community structure for each type of morphological index-dependent, single-layer brain networks and multiplex morphological brain networks at the group level. To this end, a group-level mean similarity matrix was first obtained for each type of single-layer morphological brain networks. A nonparametric method of locally adaptive network sparsification (Foti, Hughes et al. 2011) was then used to extract a binary backbone network from each of the group-level mean similarity matrix to capture their multiscale structure. The binary backbone networks were further multiplied by the corresponding group-level mean similarity matrices in an element-wise manner to derive weighted backbone networks, which were further merged into a weighted multiplex backbone network. In the weighted multiplex backbone network, the weights of inter-layer edges were the mean similarities of corresponding edges in the multiplex morphological brain networks of all participants. Subsequent community detection was conducted on the weighted backbone networks.

#### Single-layer morphological networks

The community detection is to find a specific partition of network nodes that yields the largest modularity, *Q*, which is defined as:

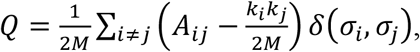

where *A*_*i j*_is the edge weight (i.e., morphological similarity) between node *i* and *j, k*_*i*_ and *k*_*j*_ are nodal strength (i.e., the sum of weights of all edges linked to a node) of node *i* and *j*, respectively, 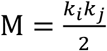 is the total weight of all edges in a network, and *σ*_*I*_ and *σ*_*j*_ represent the assignment of node *i* and *j* to a community, respectively. If node *i* and *j* are assigned to the same community, *δ*(*σ*_*i*_, *σ*_*j*_) is equal to 1, otherwise *δ*(*σ*_*i*_, *σ*_*j*_) is equal to 0. However, this method allows detection of community structure only at a single scale, and fails to identify communities smaller than the scale. Accordingly, Reichardt and colleagues (Reichardt and Bornholdt 2006) introduced a resolution parameter γ, which enables us to identify communities at multiple resolutions:

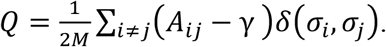

Greater γ ends up smaller modules and vice versa. In this study, the initial range of γ was set from 0.01 to 0.5 with an increment of 0.01 for each group-level, single-layer, weighted morphological backbone network. First, at each γ, we performed community detection 1,000 times and obtained a consensus matrix (Lancichinetti and Fortunato 2012) with elements indicating the proportions of each pair of nodes that were assigned to the same community over the 1,000 iterations. If there is any pair of nodes that was not assigned to the same community over the 1,000 iterations, community detection was further conducted on the consensus matrix (1,000 times) to derive a new consensus matrix. Such procedures were repeated until a consensus partition was obtained. To determine a single γ for further analysis, we then adopted a method from (He, Lim et al. 2017) to evaluate variability of the partition over different γ. Specifically, we separately calculated the variation of information (Meilă 2003) in the consensus partition for each γ with its two adjacent neighbors. The mean of the resultant two values of variation of information indicates the stability of the partition when γ changes slightly. Finally, we searched the broadest, continuous γ range with the variation of information equal to the minimum, in which the first γ was used for further analysis since the corresponding partition was stable across different resolutions. In this study, community detection of the single-layer morphological networks was performed using a spectral optimization algorithm (Newman 2006).

#### Multiplex morphological network

The modularity, *Q* for a multiplex network is defined as (Mucha 2010):

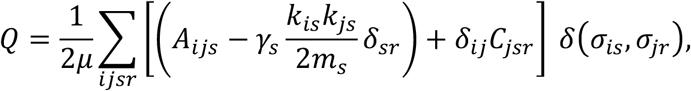

where *i* and *j* represent nodes, *s* and *r* represent layers, *A*_*ijs*_ is the edge weight (i.e., morphological similarity) between node *i* and *j* in layer *s, γ*_*s*_ is the resolution parameter in layer *s* (a constant across layers in this study), *k*_*is*_ and *k*_*js*_ are nodal strength of node *i* and *j* in layer *s*, respectively, *m*_*s*_ is the total weight of all edges in layer *s, C*_*jsr*_ is inter-layer edge weight for node *j* between layer *r* and layers, 2*μ* = ∑_*jr*_ *k*_*js*_ + *c*_*js*_ of which *k*_*js*_ = ∑_*i*_*A*_*ijs*_ denotes nodal strength of node *j* in layer *s*, and *c*_*js*_ = ∑_*r*_ *C*_*jsr*_ means total inter-layer edge weight for node *j* between layer *r* and all other layers, and *σ*_*is*_ and *σ*_*jr*_ represent the assignment of node *i* in layer *s* and *j* in layer *r* to a community, respectively. Analogous to the community detection of single-layer morphologica brain networks, a consensus partition was obtained at each γ from 0.01 to 0.50 with an increment of 0.01, and a single γ was finally determined in terms of the variation of information as a function of γ. In this study, community detection of the multiplex morphological network was performed using a heuristic method for large networks (Blondel, Guillaume et al. 2008).

### Intra- and inter-module morphological connectivity

We extracted all morphological connections within each module and between each pair of modules from each individual morphological similarity matrix (single-layer and multiple) according to the corresponding community structure identified above. The module-related morphological connections were standardized across participants (Z score transformation) and used in turn for the analyses of behavioral and cognitive association and prediction, and individual identification.

### Behavioral and cognitive association

The HCP dataset (unrelated participants) was used to examine behavioral and cognitive association with morphological brain networks using a multivariate variance component model (Ge, Reuter et al. 2016, Liegeois, Li et al. 2019). Assume that behavioral and cognitive variance is the sum of morphological connectivity effect B and environmental effect E, then the multivariate model is:

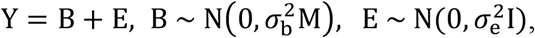

where Y is a *N*_*subject*_ *× P* _*behaviour − cognition domain*_ matrix including the behavioral and cognitive data of all participants, 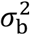 and 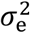 represent the variance of morphological connectivity and environment, respectively, M is an inter-individual correlation matrix of morphological connectivity, and I is an identity matrix. The multivariate model follows distributional assumptions of vec(B) ∼ N(0, ∑_B_ ⊗ M) and vec(E) ∼ N(0, ∑_E_ ⊗ I), where vec(.) is the matrix vectorization operator that converts a matrix into a vector, ⊗ denotes the Kronecker product of matrices, and Σ_*B*_ and Σ_*E*_ are *P × P* matrices estimated from M and Y. The behavioral and cognitive variance explained by morphological connectivity is evaluated as:

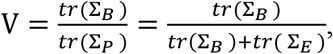

where Σ_*P*_ is the behavioral and cognitive covariance matrix, *tr*(.) is the trace operator of a matrix, and V indicates how much interindividual behavioral and cognitive variance is explained by interindividual morphological connectivity variance. These analyses were performed with publicly available codes (https://github.com/RaphaelLiegeois/FC-Behavior/). The same analyses were also performed on the LBCMLPC dataset for general validation of reproducibility

### Behavioral and cognitive prediction

Based on the same datasets as used in analyzing behavioral and cognitive association, a brain basis set modeling method (BBS) was employed to explore whether morphological brain networks are predictive of individual behavioral and cognitive performance. The BBS is a multivariate method that produces a set of components for phenotypic prediction after dimensionality reduction via principal component analysis (Sripada, Angstadt et al. 2019). To assess the performance of the BBS model, a 10-fold cross-validation was used. That is, all participants were divided into 10 non-overlapping, equally-sized subgroups, each of which was used as test dataset in turn with the others as training dataset. Specifically, in each fold, the principal component analysis was first conducted on the training dataset (*N*_*subject*_ *× P*_*morphological connectivity*_) for the dimensionality reduction. The number of retained components was determined to ensure that more than 80% variance of the training dataset were explained. Based on the expression scores of the retained components, a linear regression model was fitted between these expression scores and each behavioral and cognitive domain in the training dataset. To predict behavioral and cognitive performance for a given participant in the test dataset, the expression scores of his or her morphological connectivity in the retained components were first calculated. The predicted values were then calculated as the Dot Product between the expression scores and the fitted coefficients (i.e., beta values) for each behavior and cognitive domain derived from the training dataset. Finally, we calculated the Pearson correlation between the predicted values and real scores for each behavior and cognitive domain.

### Individual identification

The BNU dataset was used to explore whether morphological brain networks can serve as connectome fingerprints to identify individuals using the method from (Finn, Shen et al. 2015). Briefly, participants in data session 1 were first assigned as target set and participants in data session 2 as database set. For each participant in the target set, we then calculated his or her similarity (Spearman correlation) with each participant in the database set with respect to their vectorized morphological connectivity profiles. For a given participant in the target set, we regarded the identification as hitting if the largest similarity was observed for the same participant in the database set. Finally, the identification accuracy was calculated as the ratio of the number of hitting to the number of participants. These procedures were repeated by assigning participants in data session 2 as target set and data session 1 as database set. The same analyses were also performed on the 8 pairs of twins completing two structural image scans in the HCP dataset, and the LBCMLPC dataset for validation of reproducibility.

### Heritability

The HCP dataset (twin participants) was used to investigate the extent to which morphological brain networks are genetically controlled. Theoretically, MZ twins have identical genes while DZ twins share on average 50% genes, and both MZ and DZ twins share 100% common environment. In a ACE model, the variance of a phenotypic variable is assumed to be composed of additive genetic component (A), common environment (C), and unique environment (E) wherein A denotes the influences of multiple alleles at different loci on the genome on the phenotype, C denotes the same family environment, and E includes the unique living environment and measurement errors. Formally, the narrow-sense heritability is defined as the proportion of phenotypic variance that is attributed to genetic factors:

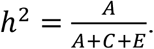

In this study, the heritability was estimated for morphological connectivity between each pair of regions for each type of single-layer morphological brain networks with the APACE package (Chen, Formisano et al. 2019).

## Statistical analysis

### Performance of behavioral and cognitive association and prediction

To test the performance of behavioral and cognitive association, we performed nonparametric permutation tests to examine whether the observed proportions of behavioral and cognitive variance explained by morphological brain networks could occur by chance. Specifically, we randomly shuffled real behavioral and cognitive scores of all domains over participants, and reran the multivariate variance component model (10,000 times). These procedures resulted in a set of null distributions, based on which a *P* value was obtained for the association analysis between each behavioral and cognitive domain and morphological connectivity within each module and between each pair of modules. A false discovery rate (FDR) procedure was used to corrected for multiple comparisons at the level of q = 0.05. Analogously, for the analysis of behavioral and cognitive prediction, we shuffled randomly real behavioral and cognitive scores of each domain over participants, reran the BBS method (10-fold cross-validation), and recomputed the correlation between predicted behavioral and cognitive values and the permutated true scores. After repeating these procedures 10,000 times, a *P* value was obtained for the prediction analysis between each behavioral and cognitive domain by morphological connectivity within each module and between each pair of modules, followed by a FDR-based multiple comparison correction.

### Significance of heritability

To locate morphological connectivity showing significant heritability, we shuffled randomly the labels (MZ or DZ) of each pair of twins (1,000 times), and recomputed the connectional heritability for each type of single-layer morphological brain networks. Based on the resultant empirical null distributions, a threshold-free network-based statistics approach (Baggio, Abos et al. 2018) was used to search morphological connections that exhibited significant heritability after correcting for family-wise error rates.

### Effects of morphological network type, morphological connectivity type, and behavioral and cognitive domain

For the proportions of explained variance from analysis of behavior and cognitive association and correlation coefficients from analysis of behavior and cognitive prediction, nonparametric Scheirer Ray Hare tests were used to examine whether they were dependent on the type of morphological connectivity (intra-module versus inter-module) and differed among different behavioral and cognitive domains. For the accuracies from analysis of individual identification, nonparametric two-tailed Wilcoxon rank sum tests were used to examine the effects of morphological connectivity type (intra-module versus inter-module). Finally, for edge-wise heritability, nonparametric kruskalwallis test was used to examine the differences among different types of single-layer morphological brain networks (CTNs versus FDNs versus GINs versus SDNs), and nonparametric two-tailed Wilcoxon rank sum tests were used to examine the effects of morphological connectivity type (intra-module versus inter-module) for each type of single-layer morphological brain networks. The FDR procedure was used to correct for multiple comparisons at the level of q = 0.05.

### Multiplex versus single-layer morphological brain networks

We examined the differences between multiplex morphological brain networks and each type of single-layer morphological brain networks in the proportions of explained variance from analysis of behavior and cognitive association, correlation coefficients from analysis of behavior and cognitive prediction, and accuracies from analysis of individual identification using nonparametric two-tailed Wilcoxon rank sum tests. The FDR procedure was used to correct for multiple comparisons at the level of q = 0.05.

## Results

### Morphological index-dependent, single-layer morphological brain networks

Figure 2A shows the group-level mean morphological similarity matrices derived from different morphological indices. Consistent with our previous study, distinct connectivity patterns were observed among different types of morphological brain networks as characterized by low spatial Spearman rank correlation coefficients (*r* = 0.013 − 0.058) (Fig. 2B). These findings suggest that different morphological indices are largely independent and complementary to each other for mapping morphological brain networks. Thus, we utilized a multiplex network model to integrate different types of morphological brain networks for subsequent analyses.

**Figure 2.**
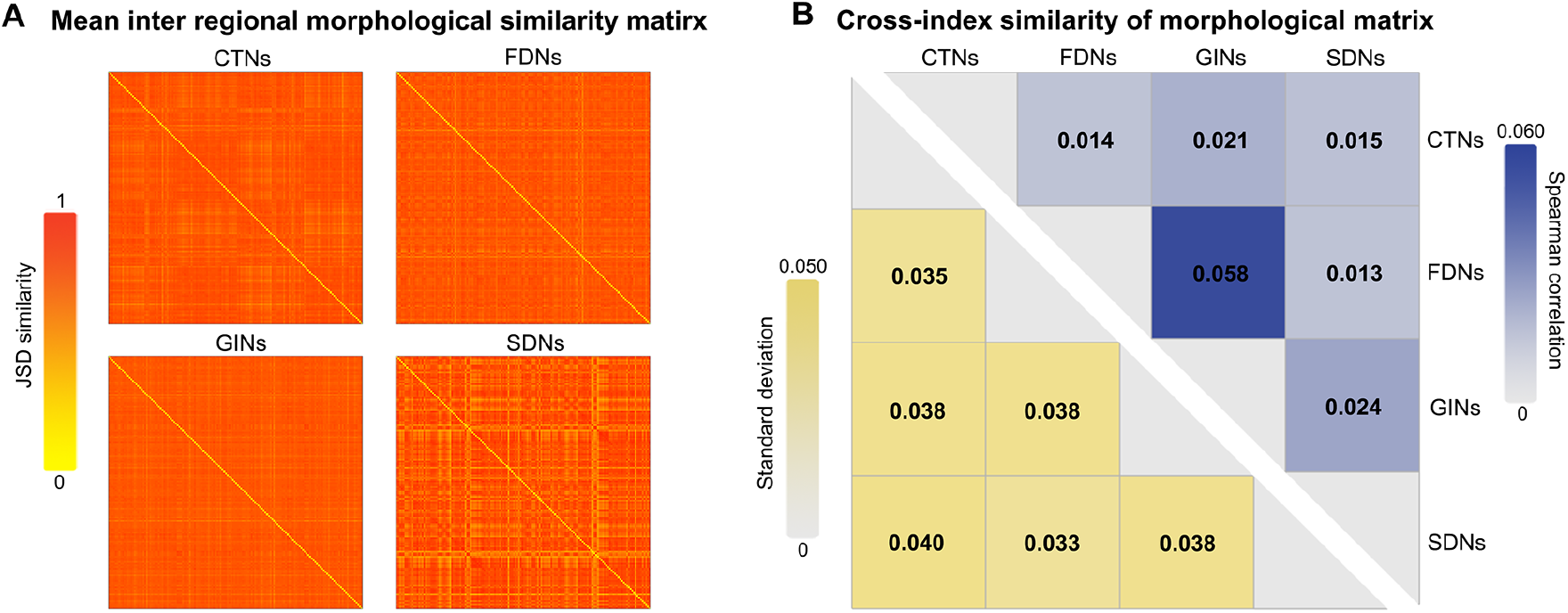
Mean interregional morphological similarity matrices and their cross-index spatial similarities (HCP dataset, unrelated participants). Although high interregional morphological similarities were observed all morphological brain networks, their spatial similarities were very low. These findings suggest distinct connectivity patterns among the morphological index-dependent, single-layer brain networks. JSD, Jensen-Shannon divergence; CTNs, cortical thickness networks; FDNs, fractal dimension networks; GINs, gyrification index networks; SDNs, sulcal depth networks.

### Community structure of multiplex morphological brain networks

Five communities or modules were identified for the group-level multiplex morphological network (*Q* = 0.402). The communities differed in size (i.e., number of regions) over the four layers with community 1 including the most regions (178 in total) and community 5 the least (43 in total). For community composition, the communities were generally composed of bilaterally symmetrical regions in each layer with varying degrees of spatial overlapping between layers (Fig. 3, and Tables S3 and S4). For example, the most stable community across layers was community 4 that mainly embraced frontal and temporal cortex, precentral gyrus and postcentral gyrus, of which nearly half were common between any pair of layers (dice coefficient = 0.415 ± 0.098). In contrast, community 2 involved widespread regions in the cortex that varied largely across layers, resulting in a small spatial overlapping between different types (dice coefficient = 0.176 ± 0.162). It should be noted that even for the most stable community 4, there were only 4 regions that were shared by all layers.

**Figure 3.**
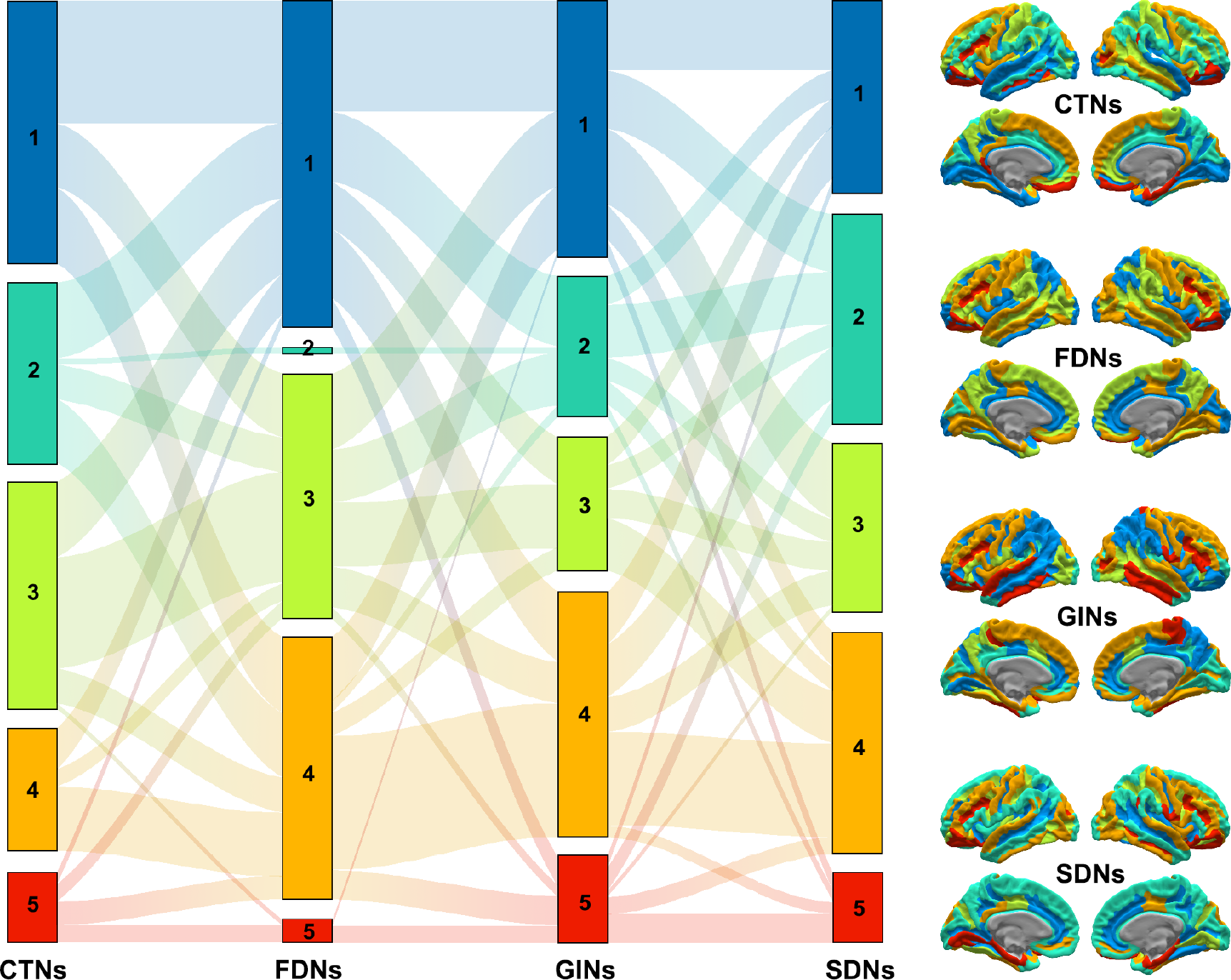
Sankey diagram and cortical surface mapping of community structure derived from group-level multiplex morphological brain network (HCP dataset, unrelated participants). The same colors correspond to the same modules between the sankey diagram and cortical surface mapping. CTNs, cortical thickness networks; FDNs, fractal dimension networks; GINs, gyrification index networks; SDNs, sulcal depth networks.

These findings further highlight distinct architecture among different types of morphological brain networks at a finer scale of community composition in addition to overall connectivity patterns.

### Multiplex morphological brain networks can explain interindividual variance in behavior and cognition

Figure 4B presents the extents to which behavioral and cognitive variance are explained by intra- and inter-module morphological connectivity. Visual inspection showed that the proportions of explained variance varied across behavioral and cognitive domains and depended on the category of morphological connectivity. Specifically, it seemed that inter-module morphological connectivity explained more behavioral and cognitive variance than intra-module morphological connectivity, and variance in the Cognition and Motor domains were accounted for to a greater degree than the other four domains. These findings were validated by further statistical analyses: 1) inter-module morphological connectivity encoded more behavioral and cognitive information than intra-module morphological connectivity (*P* < 0.001; Fig. 4C); and 2) different behavior and cognitive domains were encoded by morphological connectivity to varying degrees (*P* ∼ 0) with a pattern of Cognition > Motor > Emotion > Sensory > Personality > Alertness (*P* < 0.05, FDR corrected except for Motor vs Emotion, Emotion vs Sensory, and Personality vs Alertness) (Fig. 4D).

**Figure 4.**
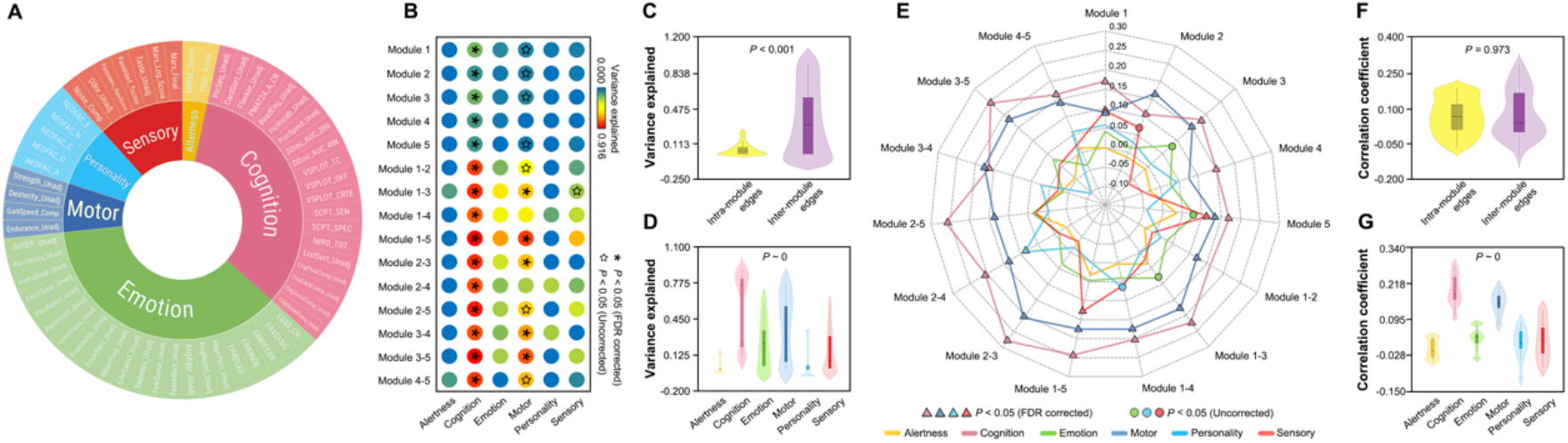
Behavioral and cognitive association and prediction (HCP dataset, unrelated participants). **A**) Sunburst plot summarizing the items included in the six behavioral and cognitive domains used in this study. **B**) The proportions of variance explained in each behavioral and cognitive domain by intra- and inter-module morphological connectivity. **C**) Violin plots showing different abilities between intra- and inter-module morphological connectivity in explaining behavioral and cognitive variance. **D**) Violin plots showing varying proportions of variance explained in different behavioral and cognitive domains by morphological connectivity. **E**) Radar chart showing the Pearson correlations between real scores and predicted values of each behavioral and cognitive domain by morphological connectivity within each module and between each pair of modules. **F**) Violin plots showing different abilities between intra- and inter-module morphological connectivity in predicting behavioral and cognitive performance. **G**) Violin plots showing varying extents among different behavioral and cognitive domains that were predictive of morphological connectivity. FDR, false discovery rate.

### Multiplex morphological brain networks can predict individual behavior and cognition

Figure 4E shows the results of BBS-based predictive models for each behavioral and cognitive domain. Significantly positive correlations were found between predicted values and actual scores for the Cognition (*r* = 0.108 − 0.280) and Motor (*r* = 0.076 − 0.206) domains when using morphological connectivity within each module (except for module 4 for the Motor domain) and inter-module morphological connectivity between each pair of modules (*P* < 0.05, FDR corrected). In addition, the Sensory domain can be predicted by morphological connectivity within module 1 and 5, and between module 1 and 5 (*r* = 0.094, 0.113 and 0.128, respectively), and the Personality domain can be predicted by morphological connectivity between module 2 and 4 (*r* = 0.088) (*P* < 0.05, FDR corrected). No significant correlations were found for the Alertness or Emotion domain (*P* > 0.05, FDR corrected). Further statistical analyses revealed that the correlation coefficients were comparable when using intra-module and inter-module morphological connectivity as features (*P* = 0.973; Fig. 4F), but differed significantly among different behavioral and cognitive domains (*P* ∼ 0), with a pattern of Cognition > Motor > Emotion > Alertness, and Cognition > Motor > Sensory and Personality (*P* < 0.05, FDR corrected; Fig. 4G).

### Multiplex morphological brain networks can distinguish individuals from each other

For the BNU dataset, low correlations were observed again between different types of single-layer morphological brain networks (*r* = 0.013 to 0.113; Fig. S1), based on which the resultant group-level multiplex morphological brain network also exhibited optimized community structure (Fig. S2). Using morphological connectivity within each module and between each pair of modules as features, almost all individuals can be successfully identified, no matter which data session was treated as target set (accuracy = 96.5% ∼ 100%; Fig. 5). Moreover, the accuracies were independent on the type of morphological connectivity (session 1 as target set: *P* = 0.330; session 2 as target set: *P* = 0.191). Interestingly, based on the 8 pairs of twin subjects in the HCP dataset who completed two structural image scans, 100% accuracies were achieved regardless of intra- or inter-module morphological connectivity.

**Figure 5.**
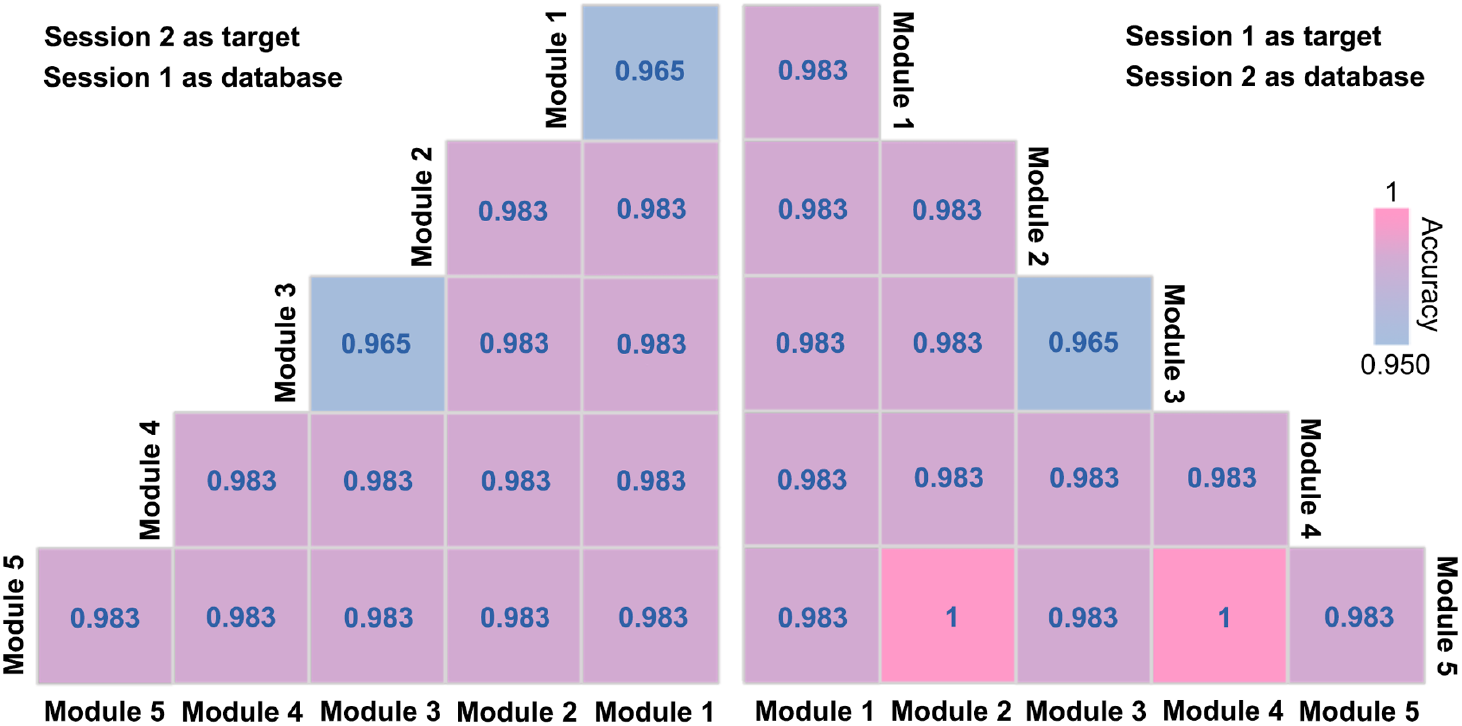
Accuracy of individual identification (BNU dataset). High accuracies were observed for morphological connectivity within each module and between each pair of modules regardless of which data session was assigned as the target set.

### Heritability of morphological brain networks

Generally, interregional morphological connectivity exhibited low-moderate heritability for each type of single-layer morphological brain networks (Fig. S3). Nevertheless, different types of single-layer morphological brain networks were associated with different levels of heritability (*P* ∼ 0), with a pattern of SDNs > FDNs > CTNs > GINs (*P* < 0.05, FDR corrected). Moreover, for each type of single-layer morphological brain networks, inter-module morphological connections exhibited significantly higher heritability than intra-module connections (*P* < 0.05, FDR corrected). Using a threshold-free network-based statistics approach, morphological connections that exhibited significant heritability were further located for each type of single-layer morphological brain networks (*P* < 0.05, FWE corrected; Fig. 6). We found that heritable morphological connections were mainly in the SDNs (280, *h*^*2*^ = 0.340 ∼ 0.637), followed by FDNs (101, *h*^*2*^ = 0.304 ∼ 0.421), CTNs (13, *h*^*2*^ = 0.380 ∼ 0.469) and GINs (6, *h*^*2*^ = 0.336 ∼ 0.397). Interestingly, the heritable morphological connections were mainly inter-module edges (SDNs: 248, 88.6%; FDNs: 74, 73.3%; CTNs: 13, 100%; GINs: 5, 83.3%).

**Figure 6.**
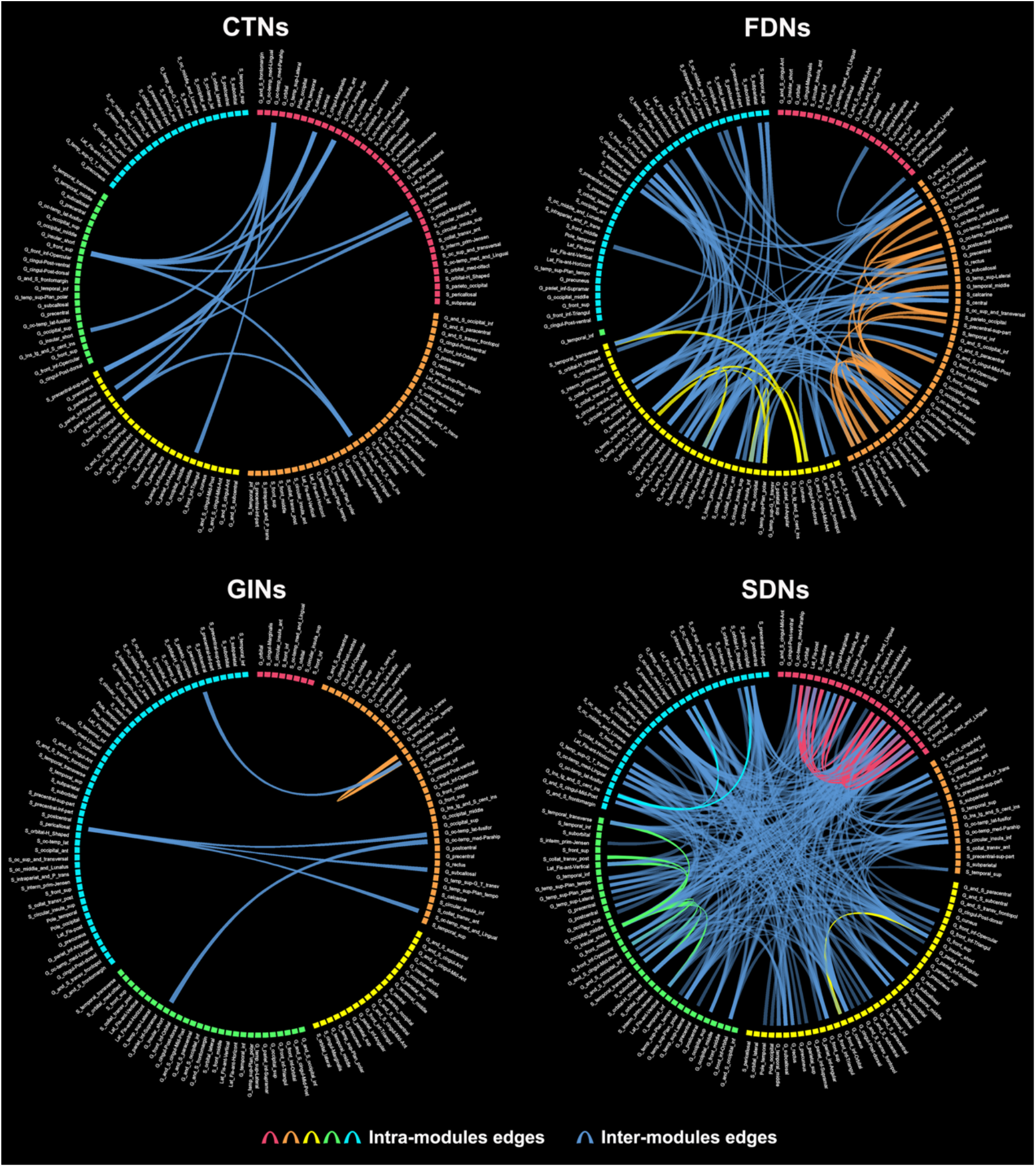
Morphological connectivity with significant heritability for each type of single-layer morphological brain networks (HCP dataset, twin participants). Brain regions share the same color if they are assigned to the same community. The line width is proportional to the heritability of corresponding connections. CTNs, cortical thickness networks; FDNs, fractal dimension networks; GINs, gyrification index networks; SDNs, sulcal depth networks.

### Differences between multiplex and single-layer morphological brain networks

As shown in Figure 7, the multiplex morphological brain networks 1) accounted for more variance in the Cognition, Emotion, Motor and Sensory domains than each type of single-layer morphological brain networks (*P* < 0.05, FDR corrected); 2) gave rise to greater correlation coefficients between predicted values and actual scores in the Cognition domain than single-layer CTNs, FDNs and GINs, and in the Motor domain than single-layer FDNs and GINs (*P* < 0.05, FDR corrected); and 3) generated higher accuracies for individual identification than single-layer CTNs, GINs and SDNs (*P* < 0.05, FDR corrected). In addition, we noted that the multiplex morphological brain networks explained less variance in the Alertness and Personality domains than all types of single-layer morphological brain networks (*P* < 0.05, FDR corrected).

**Figure 7.**
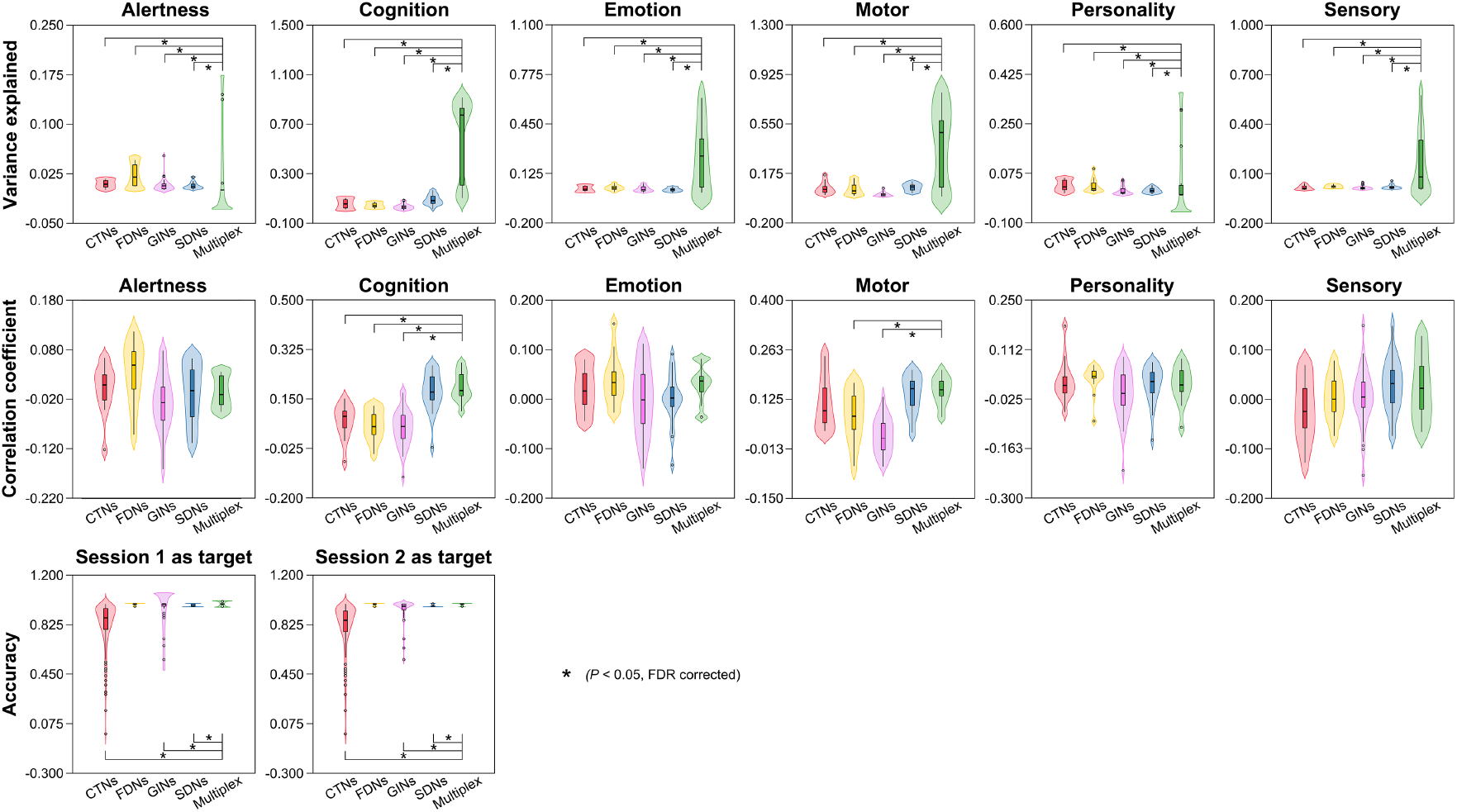
Multiplex versus single-layer morphological brain networks in behavioral and cognitive association, behavioral and cognitive prediction, and individual identification (HCP dataset, unrelated participants, and BNU dataset). Generally, compared with single-layer morphological brain networks, multiplex morphological brain networks explained more variance in behavior and cognition, generated greater correlations for behavioral and cognitive prediction, and showed higher accuracies for individual identification. CTNs, thickness network, FDNs, fractal dimension networks, GINs, gyrification index networks, SDNs, sulcal depth networks; *, *P* < 0.05, corrected by the false discovery rate procedure.

### Reproducibility of main results

Our main results were replicated on an independent LBCMLPC dataset as characterized by 1) low spatial correlations in the connectivity patterns among different types of single-layer morphological brain networks (*r* = 0.008 − 0.057; Fig. S4); 2) optimized community structure with composition varying across different types (Fig. S5); 3) morphological connectivity type-dependent and domain-specific behavioral and cognitive association (*P* ∼ 0 and *P* = 0.014, respectively; Fig. S6B-D) and prediction (both *P* < 0.001; Fig. S6E-G); 4) high accuracies for individual identification (81.6% − 100%; Fig. S7); and 5) more explained variance, greater correlations and higher accuracies for multiplex than single-layer morphological brain networks in the analyses of behavioral and cognitive association, behavioral and cognitive prediction, and individual identification, respectively (*P* < 0.05, FDR corrected; Fig. S8).

## Discussion

In this study, we investigated behavioral and cognitive implications, potential as fingerprints, and genetic basis of individual-level morphological brain networks. We found that morphological brain networks were associated with and predictive of multiple behavioral and cognitive domains, capable of recognizing individuals even twin subjects with high accuracies, and under genetic control. Moreover, the findings were largely reproducible across different datasets. Collectively, these findings deepen our understanding of interindividual differences in morphological brain networks and provide strong evidence for the usage of individual-level morphological brain networks in future brain connectome research.

### Behavioral and cognitive relevance of morphological brain networks

By associating interindividual differences in brain structure and function with various behavior and cognition, accumulating evidence indicates that human brain networks are physiological basis for cognitive performance and a diverse repertoire of behaviors (Bullmore and Sporns 2009). Currently, behavioral and cognitive relevance of structural and functional brain networks has been well documented in the literature in health and disease (Siegel, Ramsey et al. 2016, Liegeois, Li et al. 2019, Sripada, Angstadt et al. 2019, Lee, Rodrigue et al. 2020, Lin, Baete et al. 2020, Tian, Margulies et al. 2020). However, relevant studies are rare for morphological brain networks (Tijms, Yeung et al. 2014, Seidlitz, Vasa et al. 2018). Combining multiple behavioral and cognitive domains with multivariate approaches, here we showed that individual-level morphological brain networks can capture interindividual differences in multiple domains. In particular, the Cognition and Motor domains were predicted by morphological connectivity within each module and between any pair of modules, indicating general involvement of whole-brain morphological connectivity in these two domains. In contrast, the Personality and Sensory domains were only predicted by morphological connectivity of specific modules, suggesting selective implication of a specific set of morphological connectivity in these two domains. Overall, these findings suggest that morphological brain networks can function as potential biomarkers for individual performance in the four domains. However, the Alertness and Emotion domains cannot be predicted by morphological brain networks, which may be not sensitive to the features or models used in this study. Interestingly, we found that inter-module morphological connectivity encoded more interindividual variance in behavior and cognition than intra-module morphological connectivity. This is consistent with previous findings from meta-analysis of functional co-activation patterns showing that brain regions with many inter-module connections increased activity in tasks that engaged in multiple cognitive functions and exhibited a similar spatial distribution to areas associated with many cognitive functions (Bertolero, Yeo et al. 2015). Furthermore, evidence from meta-analysis of functional connectivity in eight psychiatric disorders revealed that inter-network connectivity alterations were shared in the disorders and the connectivity alterations were related to general cognitive performance (Sha, Xia et al. 2018). These structural and functional findings collectively suggest vital roles of inter-module connections in promoting behavioral and cognitive performance. Future studies are required to explore whether inter-module connections could, to a greater extent, determine behavioral and cognitive performance in health and disease by combining different modalities of brain networks.

### Individual identification through morphological brain networks

Recently, more and more studies have shown that functional connectivity profiles can act as connectome fingerprints to accurately identify subjects (Finn, Shen et al. 2015, Miranda-Dominguez, Feczko et al. 2018, Byrge and Kennedy 2019, Horien, Shen et al. 2019, Demeter, Engelhardt et al. 2020). Moreover, this type of connectome fingerprints is robust regardless of whether the brain is at rest or engaged in external stimuli during imaging (Finn, Shen et al. 2015), and is similar between youths and adults (Demeter, Engelhardt et al. 2020). Here, we extended connectome fingerprints from functional connectivity profiles to morphological brain networks by demonstrating that interregional morphological connectivity patterns can also identify individuals with high accuracies even for twin subjects. These findings suggest that individual-level morphological brain networks can serve as a new type of connectome fingerprints for individual identification. Nevertheless, we noted two marked differences between functional and morphological connectome fingerprints for individual identification. First, previous functional studies consistently showed that the default-mode and frontoparietal networks were the most highly distinctive among individuals (Finn, Shen et al. 2015, Miranda-Dominguez, Feczko et al. 2018, Horien, Shen et al. 2019, Demeter, Engelhardt et al. 2020). However, in this study we found that morphological connectivity within each module and between each pair of modules exhibited high and comparable accuracies in identifying individuals. This is consistent with a recent functional network study showing that individually distinct information was widespread throughout the cortex and no specific connections or networks were required for individual identification (Byrge and Kennedy 2019). The other difference was that in contrast to low accuracies in identifying twin subjects based on functional connectivity profiles (accuracy < 60%; (Demeter, Engelhardt et al. 2020), morphological brain networks distinguished twins from each other with 100% accuracies. The discrepancy may reflect different degrees and ranges of genetic effects on functional and morphological brain networks, which is an interesting topic for future studies. Notably, due to the small sample size for the twin participants used for individual identification, further studies with more twins are required to test the reproducibility of our findings. Overall, given the excellent, reproducible performance of morphological brain networks in distinguish individuals, together with their high test-retest reliability as demonstrated in our previous study (Li et al., 2020), we propose this relatively new type of brain networks as an important tool for individualized studies in cognitive and clinical neuroscience.

### Genetic basis of morphological brain networks

Imaging genetics, which refers to the use of neuroimaging techniques to assess the effects of genetic factors on brain structure and function, has significantly advanced our understanding of genetic architecture of the human brain that are relevant to brain disorders, development and aging (Jansen, Mous et al. 2015, Maggioni, Squarcina et al. 2020). In the context of large-scale brain networks, genetic associations have been relatively well-established for structural and functional brain networks (Colclough, Smith et al. 2017, Sudre, Choudhuri et al. 2017, Elliott, Sharp et al. 2018, Zhong, Wei et al. 2020). For morphological brain networks, two previous human studies found that they shared similar topological properties to a transcriptional brain network (Romero-Garcia, Whitaker et al. 2018), and had greater interregional morphological connectivity for regions with stronger gene coexpression (Seidlitz, Vasa et al. 2018). Similar findings were also observed in mouse (Yee, Fernandes et al. 2018). These findings collectively indicate key roles of genes on overall wiring patterns of morphological brain networks. Here, our results further deepen the understanding of genetic effects on morphological brain networks by mapping edge-wise heritability and demonstrating that the degree of heritability varied across different types of morphological brain networks and differed between intra- and inter-module connections. Specifically, we found that the SDNs exhibited the greatest heritability, followed by FDNs, CTNs and GINs. This pattern is largely comparable with differences in test-retest reliabilities among the four types of morphological brain networks (Li et al., 2020). Presumably, this may be because greater heritability is associated with smaller within-subject variance due to less effects of environment, which leads to higher test-retest reliability. Interestingly, despite different degrees of heritability among the four types of morphological brain networks, inter-module connections were consistently observed to be genetically controlled to a greater extent than intra-module connections in terms of both the heritability and the number of significantly heritable connections. As we discussed above, inter-module connections are vital for brain regions to integrating multiple cognitive functions during complex tasks. Previous studies have shown that regions that correlated most strongly with cognitive abilities were also those that were highly heritable (Thompson, Cannon et al. 2001), and areas associated with more complex reasoning abilities became increasingly heritable with maturation (Lenroot and Giedd 2008). On the other hand, evidence from computational modeling studies has shown that an efficient means to facilitate brain connectivity and functional development is to increase cortical folding (Ruppin 1993, Murre 1995), which may result from different neurodevelopmental rates of expansion of superficial and deep cortical layers. A recent study found that human cortical expansion was largely determined by cortical gene expression profile relevant to human evolution, in particular for high-order functional network (Wei, de Lange et al. 2019), which are routinely composed of association cortex that integrate information from different systems to facilitate task performance via adaptively adjusting inter-network interactions (Cole, Reynolds et al. 2013). Based on these findings together with our results, we speculate that the higher heritability for inter-module connections may be the consequence of evolution, which ensures the overall cognitive performance of human brain in response to complex tasks by prohibiting an excess of modifications by environmental experience on these key connections.

### Multiplex versus single-layer morphological brain networks

We found that multiplex morphological brain networks outperformed each type of single-layer morphological brain networks in terms of the explained variance in and predictive ability of behavior and cognition, and the accuracy for individual identification. These findings suggest that brain networks derived from different morphological indices are complementary to each other in determining individual uniqueness. This is consistent with previous findings that different types of single-layer morphological brain networks exhibited low spatial similarities in their connectivity patterns (Li et al., 2020) and different sensitivities in revealing disease-related alterations (Lv et al., 2020), and that taking inter-layer similarity of morphological brain networks improved late mild cognitive impairment/Alzheimer’s disease classification (Mahjoub, Mahjoub et al. 2018). Presumably, different cellular mechanisms, distinct genetic origins (Panizzon, Fennema-Notestine et al. 2009, Winkler, Kochunov et al. 2010, Strike, Hansell et al. 2019) and differential developmental/aging trajectories (Raznahan, Shaw et al. 2011, Hogstrom, Westlye et al. 2013, Wierenga, Langen et al. 2014) of the morphological indices may account for, to some extent, the observed and reported differences among different types of morphological brain networks (notably, although surface area was not included in this study, it is directly related to cortical folding). Collectively, our findings highlight the necessity to fuse different morphological indices for a more complete mapping and understanding of morphological brain networks in typical and atypical populations.

### Limitation and future directions

First, the widely used Destrieux atlas was used in this study to define network nodes. Given accumulating evidence for brain parcellation-dependent connectivity patterns and topological organization for functional and structural as well as morphological brain networks (Wang, Wang et al. 2009, Zalesky, Fornito et al. 2010, Wang, Jin et al. 2016), it is important in the future to test whether our findings are reproducible when using different atlases for brain parcellation, in particular to determine which atlas will give rise to the best performance in accounting for and predicting individual behavior and cognition. Second, only four morphological indices were used in this study to construct multiplex brain networks. Including other morphological indexes that are computationally available from different toolboxes, such as surface area and surface normal, may help more comprehensively unveil the relationships between morphological brain networks and behavior and cognition. Third, using a population-based approach, a previous study showed poor correspondences between morphological covariance and structural/functional connectivity patterns (Reid, Lewis et al. 2016). For the JSD-based individual-level morphological brain networks, it is interesting to explore their similarities and differences with other modalities of brain networks, and to rank different modalities of brain networks in terms of their abilities to underlie behavior and cognition. Finally, all analyses in this study were conducted at the connectional level. Our previous studies have shown that the morphological connectivity was wired in an optimized manner to exhibit nontrivial network organization (Wang, Jin et al. 2016, Li et al., 2020). Future studies are warranted to explore interindividual differences in system-level behavior of morphological brain networks in the context of behavioral and cognitive relevance, individual identification and genetic basis.

---

**Figure S1.**
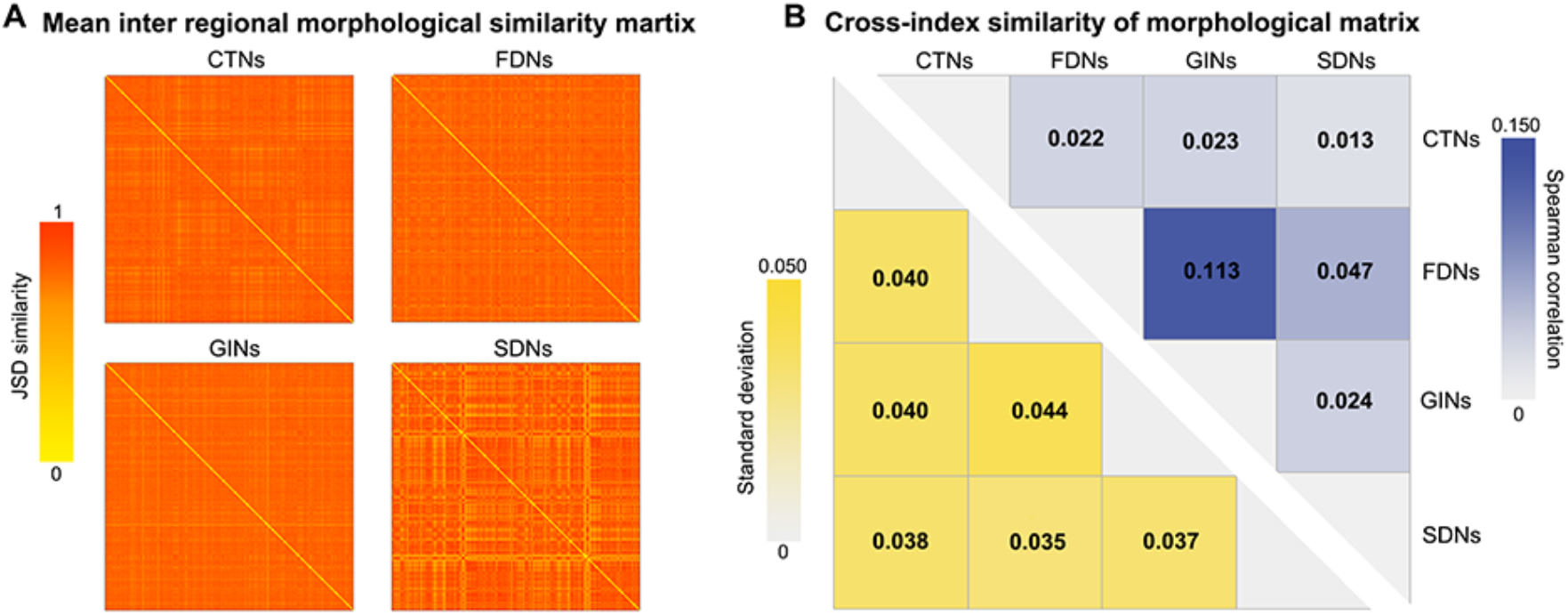
Mean interregional morphological similarity matrices and their cross-index spatial similarities (BNU dataset).

**Figure S2.**
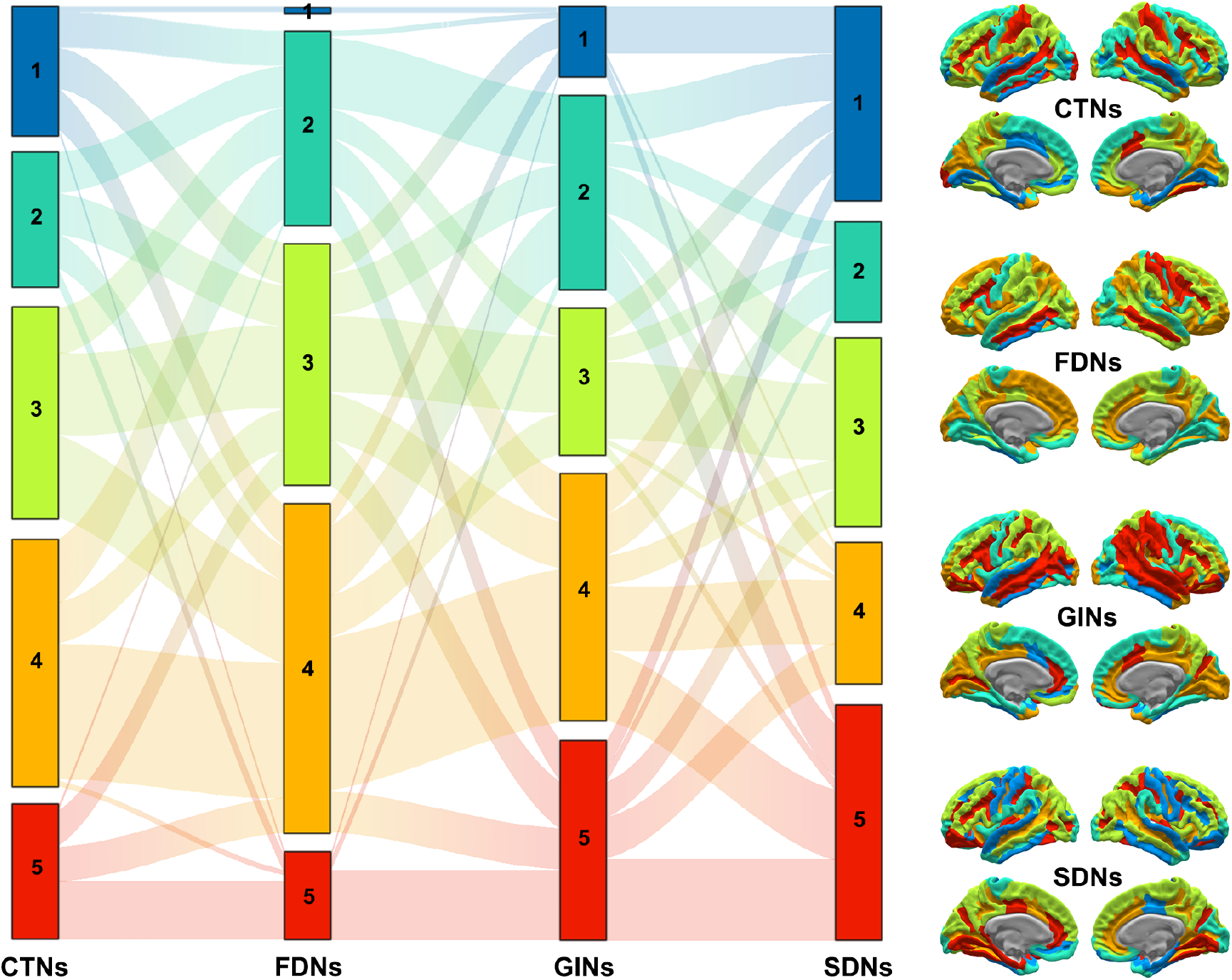
Sankey diagram and cortical surface mapping of community structure derived from group-level multiplex morphological brain network (BNU dataset).

**Figure S3.**
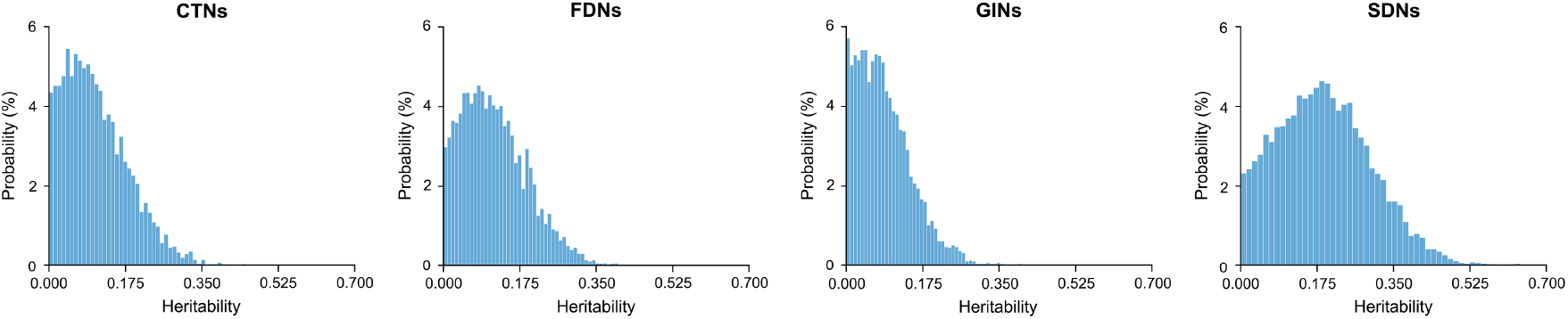
Distribution of heritability of all morphological connections for each type of single-layer morphological brain networks (HCP dataset, twin participants).

**Figure S4.**
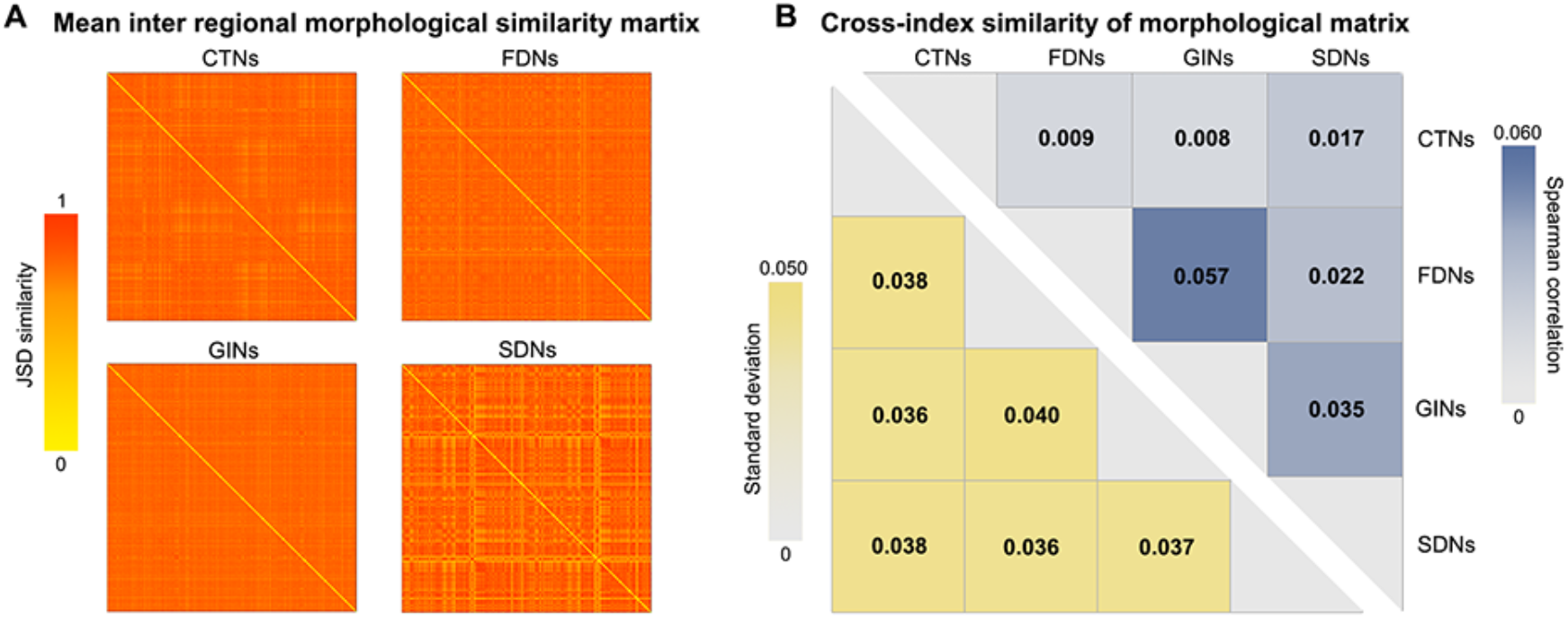
Mean interregional morphological similarity matrices and their cross-index spatial similarities (LBCMLPC dataset).

**Figure S5.**
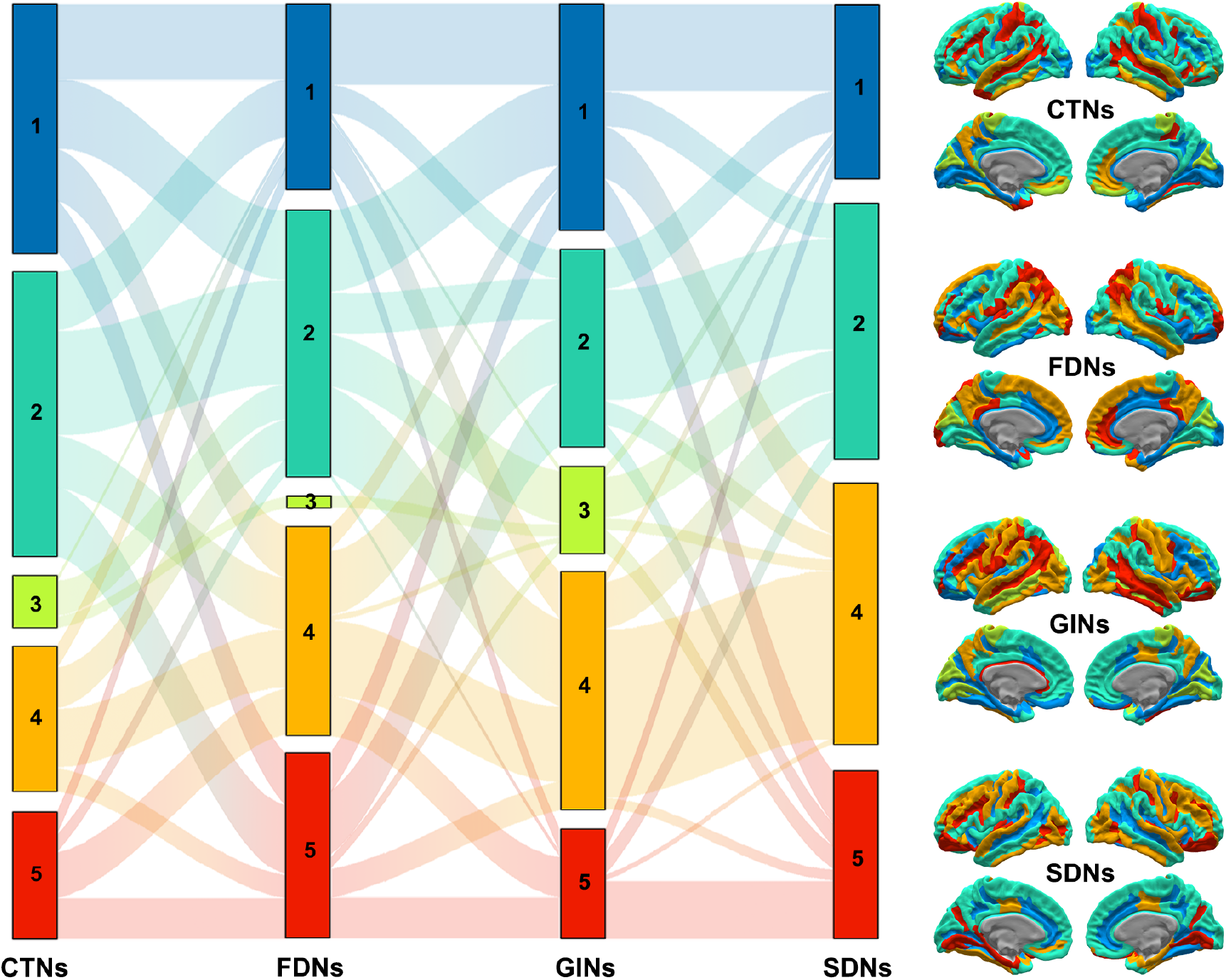
Sankey diagram and cortical surface mapping of community structure derived from group-level multiplex morphological brain network (LBCMLPC dataset).

**Figure S6.**
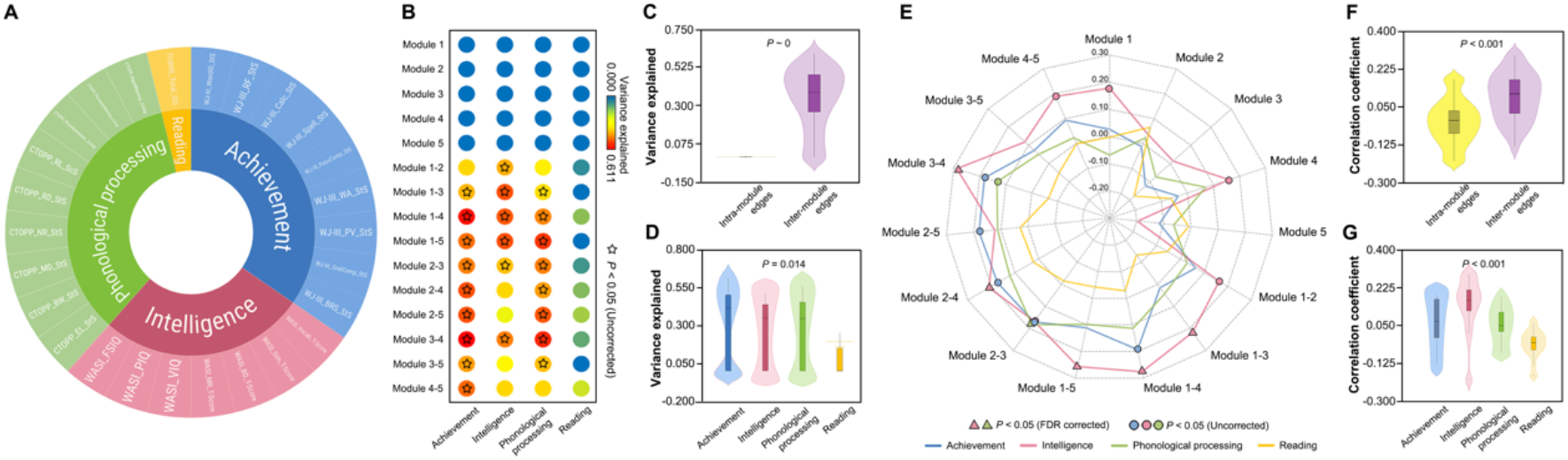
Behavioral and cognitive association and prediction (LBCMLPC dataset).

**Figure S7.**
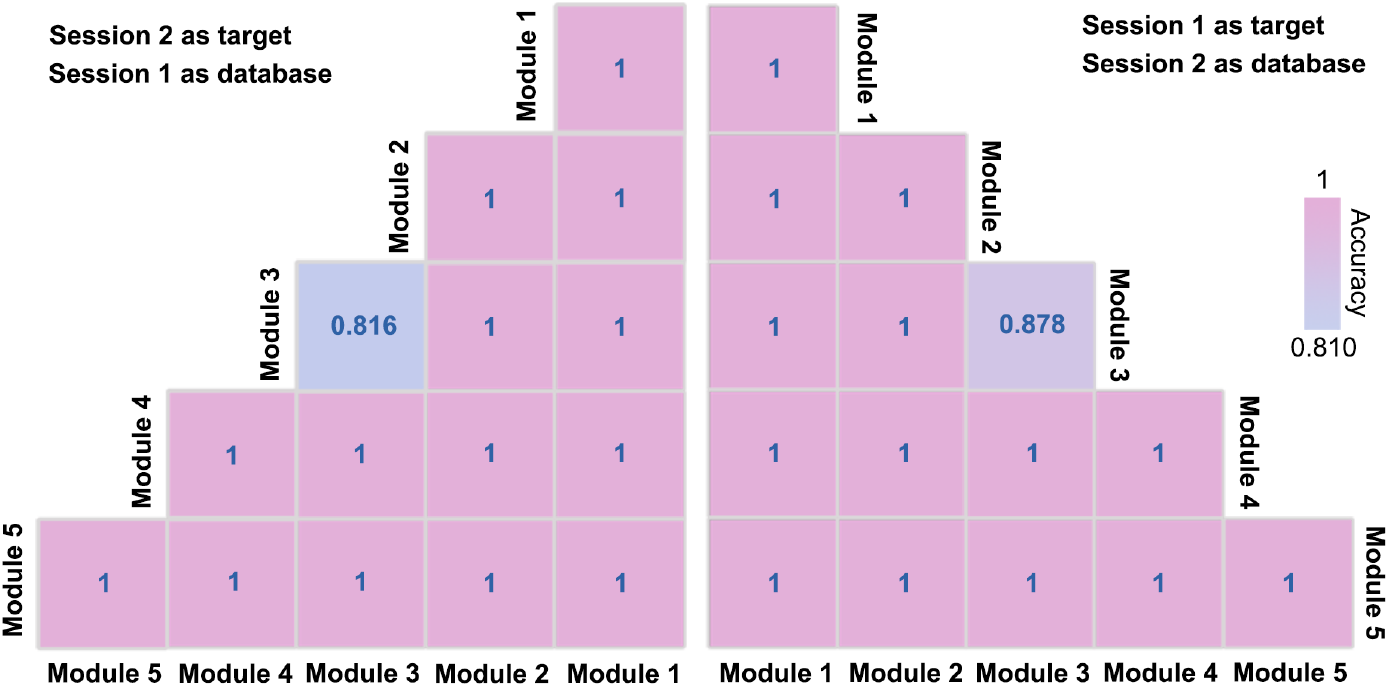
Accuracy of individual identification (LBCMLPC dataset).

**Figure S8.**
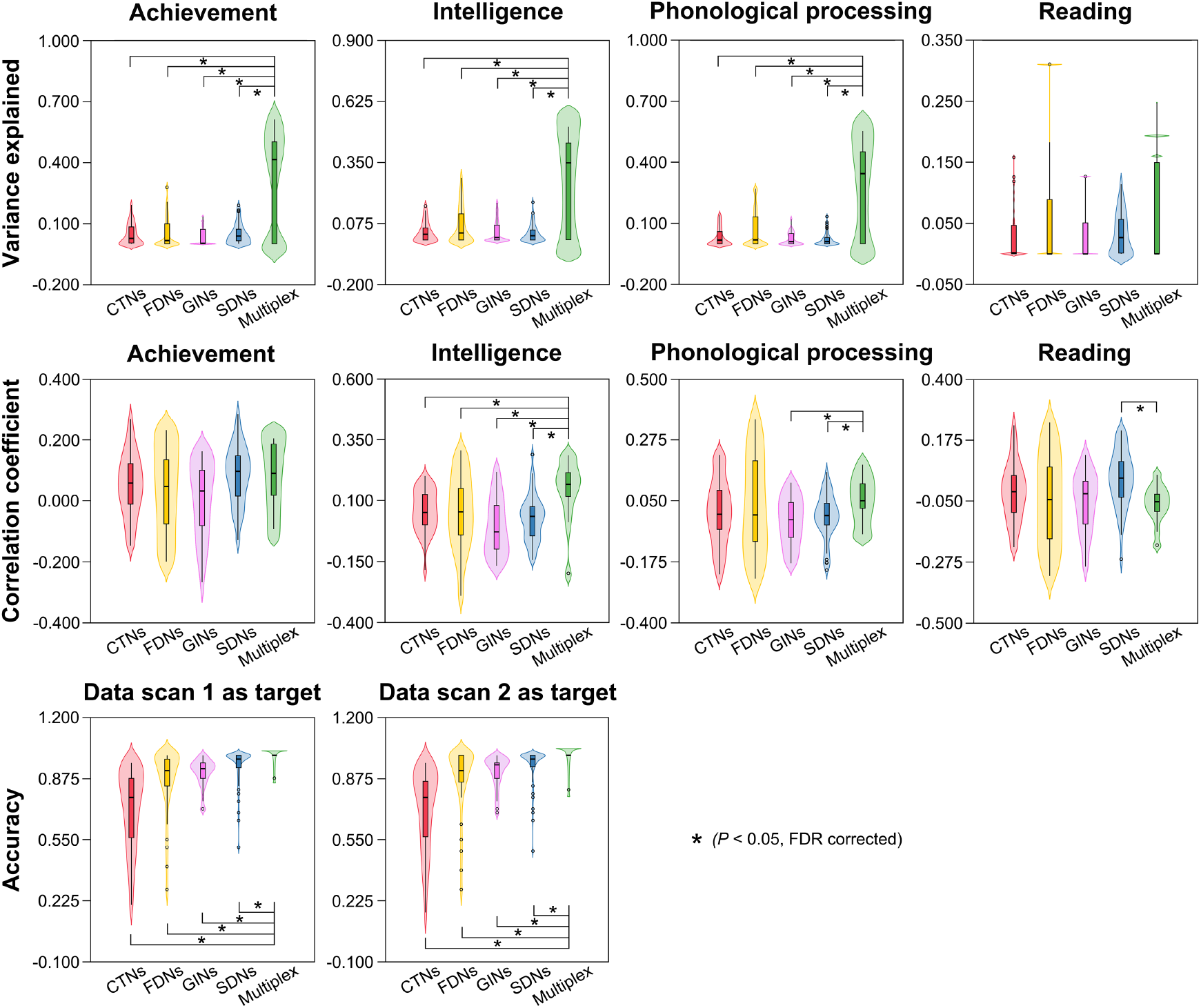
Multiplex versus single-layer morphological brain networks in behavioral and cognitive association, behavioral and cognitive prediction and individual identification (LBCMLPC dataset).

