## Supplementary material for "Towards understanding interindividual differences in cortical morphological brain networks": SI

**Behavioral and cognitive measures**

**HCP Dataset**. For the initial 581 items of behavioral measures included in the HCP dataset, we performed a multi-step screening strategy in terms of our research purpose. First, items related to health and family history, psychiatric and life function and substance use were excluded. Items resulted from MEG and fMRI tasks (e.g., accuracy and reaction time) were also deleted. For the remaining items, we further ruled out those that had the same values or were missing for more than 80% of the participants since they may fail to sufficiently capture interindividual differences. Furthermore, for items with unadjusted and adjusted values by age, original scores were used since using age-adjusted versions has little effect on the relationships between brain networks and behavior (Smith et al., 2015). In addition, if total scores were provided for several items, the total scores were retained instead of individual items. After these procedures, a total of 60 items were remained in this study and categorized into 6 behavioral and cognitive domains (Table S1). Based on the 60 items, several participants were excluded who did not complete any tests in at least one behavioral and cognitive domain from this study. Finally, to exclude effects of outliers, values that were 3.29 standard deviation (std) above and below the mean in each item were replaced by mean ± 3.29 std, respectively.

**LBCMLPC Dataset**. The LBCMLPC dataset included 50 behavioral items at data scan 1. First, if both raw scores and standardized or T-scores were available for an item, the latter were kept. Again, if total scores were available, we only retained the total scores rather than individual items. These procedures resulted in a total of 26 items left, which are categorized into 4 behavioral and cognitive domains (Table S2). No participants were excluded in terms of the criteria mentioned above. Similar to the HCP dataset, outliers were finally identified in each item and were replaced by corresponding mean ± 3.29 std.
