## Supplementary material for "Towards understanding interindividual differences in cortical morphological brain networks": Table S1

**Table S1**. Behavioral and cognitive measures included in this study for the HCP dataset (unrelated participants)

| **Domain** | **Subdomain (Test)** | **Item** |
| --- | --- | --- |
| Alertness | Cognitive Status (Mini Mental Status Examination) | MMSE_Score |
|  | Sleep (Pittsburgh Sleep Quality Index) | PSQI_Score |
| Cognition | Episodic Memory (Picture Sequence Memory) | PicSeq_Unadj |
|  | Cognitive Flexibility (Dimensional Change Card Sort) | CardSort_Unadj |
|  | Inhibition (Flanker Inhibitory Control and Attention Task) | Flanker_Unadj |
|  | Fluid Intelligence (Penn Progressive Matrices) | PMAT24_A_CR |
|  | Language/Reading Decoding (Oral Reading Recognition) | ReadEng_Unadj |
|  | Language/Vocabulary Comprehension (Picture Vocabulary) | PicVocab_Unadj |
|  | Processing Speed (Pattern Comparison Processing Speed) | ProcSpeed_Unadj |
|  | Self-regulation/Impulsivity (Delay Discounting) | DDisc_AUC_200 |
|  |  | DDisc_AUC_40K |
|  | Spatial Orientation (Variable Short Penn Line Orientation Test) | VSPLOT_TC |
|  |  | VSPLOT_OFF |
|  |  | VSPLOT_CRTE |
|  | Sustained Attention (Short Penn Continuous Performance Test) | SCPT_SEN |
|  |  | SCPT_SPEC |
|  | Verbal Episodic Memory (Penn Word Memory Test) | IWRD_TOT |
|  | Working Memory (List Sorting) | ListSort_Unadj |
|  | Cognition Summary Scores (Cognitive Function Composite Scores) | CogFluidComp_Unadj |
|  |  | CogEarlyComp_Unadj |
|  |  | CogTotalComp_Unadj |
|  |  | CogCrystalComp_Unadj |
| Emotion | Emotion Recognition (Penn Emotion Recognition Test) | ER40_CR |
|  |  | ER40ANG |
|  |  | ER40FEAR |
|  |  | ER40NOE |
|  |  | ER40SAD |
|  | Negative Affect (Self-report Questionnaire) | AngAffect_Unadj |
|  |  | AngHostil_Unadj |
|  |  | AngAggr_Unadj |
|  |  | FearAffect_Unadj |
|  |  | FearSomat_Unadj |
|  |  | Sadness_Unadj |
|  | Psychological Well-being (Self-report Questionnaire) | LifeSatisf_Unadj |
|  |  | MeanPurp_Unadj |
|  |  | PosAffect_Unadj |
|  | Social Relationships (Self-report Questionnaire) | Friendship_Unadj |
|  |  | Loneliness_Unadj |
|  |  | PercHostil_Unadj |
|  |  | PercReject_Unadj |
|  |  | EmotSupp_Unadj |
|  |  | InstruSupp_Unadj |
|  | Stress and Self-Efficacy (Self-report Questionnaire) | PercStress_Unadj |
|  |  | SelfEff_Unadj |
| Motor | Endurance (2-minute Walk Test) | Endurance_Unadj |
|  | Locomotion (4-meter Walk Test) | GaitSpeed_Comp |
|  | Dexterity (9-hole Pegboard) | Dexterity_Unadj |
|  | Strength (Grip Strength Dynamometry) | Strength_Unadj |
| Personality | Five Factor Model (NEO-FFI) | NEOFAC_A |
|  |  | NEOFAC_O |
|  |  | NEOFAC_C |
|  |  | NEOFAC_N |
|  |  | NEOFAC_E |
| Sensory | Audition (Words in Noise) | Noise_Comp |
|  | Olfaction (Odor Identification Test) | Odor_Unadj |
|  | Pain (Pain Intensity and Interference Surveys) | PainIntens_RawScore |
|  |  | PainInterf_Tscore |
|  | Taste (Regional Taste Intensity Test) | Taste_Unadj |
|  | Contrast Sensitivity (Mars Contrast Sensitivity) | Mars_Log_Score |
|  |  | Mars_Final |
