## Supplementary material for "Towards understanding interindividual differences in cortical morphological brain networks": Table S2

**Table S2**. Behavioral and cognitive measures included in this study for the LBCMLPC dataset (data scan 1)

| **Domain (Test)** | **Subdomain** | **Item** |
| --- | --- | --- |
| Achievement  (Woodcock-Johnson III) | Letter-Word Identification | WJ-III_WordID_StS |
|  | Reading Fluency | WJ-III_RF_StS |
|  | Calculation | WJ-III_Calc_StS |
|  | Spelling | WJ-III_Spell_StS |
|  | Passage Comprehension | WJ-III_PassComp_StS |
|  | Word Attack | WJ-III_WA_StS |
|  | Picture Vocabulary | WJ-III_PV_StS |
|  | Oral Comprehension | WJ-III_OralComp_StS |
|  | Basic Reading Skills | WJ-III_BRS_StS |
| Intelligence  (Wechsler Abbreviated Scale of Intelligence) | Vocabulary | WASI_Vocab_T-Score |
|  | Similarities | WASI_Sim_T-Score |
|  | Block Design | WASI_BD_T-Score |
|  | Matrix Reasoning | WASI_MR_T-Score |
|  | Verbal Intelligence | WASI_VIQ |
|  | Performance Intelligence | WASI_PIQ |
|  | Full Intelligence | WASI_FSIQ |
| Phonological Processing  (Comprehensive Test of Phonological Processing) | Elision | CTOPP_EL_StS |
|  | Blending Words | CTOPP_BW_StS |
|  | Memory for Digits | CTOPP_MD_StS |
|  | Nonword Repetition | CTOPP_NR_StS |
|  | Rapid Digit Naming | CTOPP_RD_StS |
|  | Rapid Letter Naming | CTOPP_RL_ScS |
|  | Phonemic Awareness | CTOPP_PhonAwareness _Comp |
|  | Phonemic Memory | CTOPP_PhonemicMemory_Comp |
|  | Rapid Naming | CTOPP_RapidNaming _Comp |
| Reading  (Test of Word Reading Efficiency) | Total Word Reading Efficiency | TOWRE_Total_StS |
