## Supplementary material for "Towards understanding interindividual differences in cortical morphological brain networks": Table S3

**Table S3**. Modular assignment of each region in different layers of multiplex morphological brain networks

| Region | Hemisphere | CTNs | FDNs | GINs | SDNs |
| --- | --- | --- | --- | --- | --- |
| G_and_S_frontomargi | L | 1 | 1 | 1 | 3 |
| G_and_S_occipital_inf | L | 2 | 4 | 3 | 4 |
| G_and_S_paracentral | L | 3 | 4 | 4 | 2 |
| G_and_S_subcentral | L | 2 | 1 | 3 | 2 |
| G_and_S_transv_frontopol | L | 3 | 1 | 1 | 2 |
| G_and_S_cingul-Ant | L | 2 | 1 | 1 | 1 |
| G_and_S_cingul-Mid-Ant | L | 2 | 1 | 3 | 1 |
| G_and_S_cingul-Mid-Post | L | 2 | 4 | 1 | 3 |
| G_cingul-Post-dorsal | L | 4 | 1 | 2 | 4 |
| G_cingul-Post-ventral | L | 5 | 3 | 4 | 1 |
| G_cuneus | L | 2 | 2 | 2 | 2 |
| G_front_inf-Opercular | L | 4 | 4 | 4 | 2 |
| G_front_inf-Orbital | L | 3 | 4 | 3 | 4 |
| G_front_inf-Triangul | L | 2 | 3 | 3 | 2 |
| G_front_middle | L | 2 | 4 | 4 | 4 |
| G_front_sup | L | 4 | 3 | 4 | 2 |
| G_Ins_lg_and_S_cent_ins | L | 4 | 1 | 4 | 3 |
| G_insular_short | L | 4 | 1 | 4 | 2 |
| G_occipital_middle | L | 2 | 3 | 3 | 4 |
| G_occipital_sup | L | 4 | 4 | 4 | 4 |
| G_oc-temp_lat-fusifor | L | 4 | 3 | 4 | 3 |
| G_oc-temp_med-Lingual | L | 1 | 4 | 1 | 5 |
| G_oc-temp_med-Parahip | L | 1 | 4 | 4 | 5 |
| G_orbital | L | 5 | 5 | 5 | 5 |
| G_pariet_inf-Angular | L | 2 | 1 | 1 | 2 |
| G_pariet_inf-Supramar | L | 2 | 3 | 1 | 2 |
| G_parietal_sup | L | 2 | 1 | 1 | 2 |
| G_postcentral | L | 3 | 4 | 4 | 4 |
| G_precentral | L | 4 | 4 | 4 | 4 |
| G_precuneus | L | 3 | 3 | 3 | 2 |
| G_rectus | L | 5 | 4 | 4 | 2 |
| G_subcallosal | L | 4 | 4 | 4 | 2 |
| G_temp_sup-G_T_transv | L | 3 | 1 | 4 | 3 |
| G_temp_sup-Lateral | L | 1 | 4 | 5 | 4 |
| G_temp_sup-Plan_polar | L | 4 | 1 | 4 | 4 |
| G_temp_sup-Plan_tempo | L | 3 | 3 | 4 | 4 |
| G_temporal_inf | L | 1 | 1 | 5 | 4 |
| G_temporal_middle | L | 2 | 4 | 1 | 2 |
| Lat_Fis-ant-Horizont | L | 3 | 3 | 3 | 3 |
| Lat_Fis-ant-Vertical | L | 3 | 3 | 3 | 4 |
| Lat_Fis-post | L | 3 | 2 | 1 |  |
| Pole_occipital | L | 1 | 1 | 2 | 2 |
| Pole_temporal | L | 1 | 3 | 2 | 2 |
| S_calcarine | L | 1 | 4 | 4 | 1 |
| S_central | L | 1 | 4 | 1 | 1 |
| S_cingul-Marginalis | L | 1 | 1 | 5 | 1 |
| S_circular_insula_ant | L | 1 | 3 | 1 | 1 |
| S_circular_insula_inf | L | 3 | 1 | 4 | 1 |
| S_circular_insula_sup | L | 1 | 1 | 1 | 1 |
| S_collat_transv_ant | L | 1 | 1 | 1 | 1 |
| S_collat_transv_post | L | 3 | 1 | 3 | 4 |
| S_front_inf | L | 5 | 5 | 5 | 5 |
| S_front_middle | L | 3 | 3 | 3 | 1 |
| S_front_sup | L | 3 | 1 | 1 | 3 |
| S_interm_prim-Jensen | L | 3 | 1 | 2 | 4 |
| S_intrapariet_and_P_trans | L | 3 | 1 | 1 | 1 |
| S_oc_middle_and_Lunatus | L | 3 | 3 | 3 | 3 |
| S_oc_sup_and_transversal | L | 1 | 4 | 1 | 5 |
| S_occipital_ant | L | 1 | 3 | 1 | 3 |
| S_oc-temp_lat | L | 5 | 1 | 1 | 3 |
| S_oc-temp_med_and_Lingual | L | 1 | 1 | 3 | 1 |
| S_orbital_lateral | L | 3 | 3 | 1 | 4 |
| S_orbital_med-olfact | L | 1 | 1 | 4 | 3 |
| S_orbital-H_Shaped | L | 1 | 1 | 2 | 5 |
| S_parieto_occipital | L | 1 | 4 | 1 | 3 |
| S_postcentral | L | 1 | 1 | 2 | 2 |
| S_pericallosal | L | 3 | 3 | 1 | 3 |
| S_precentral-inf-part | L | 3 | 3 | 1 | 3 |
| S_precentral-sup-part | L | 2 | 4 | 1 | 1 |
| S_suborbital | L | 3 | 3 | 2 | 4 |
| S_subparietal | L | 1 | 3 | 2 | 1 |
| S_temporal_inf | L | 5 | 4 | 5 | 2 |
| S_temporal_sup | L | 1 | 3 | 1 | 1 |
| S_temporal_transverse | L | 1 | 1 | 3 | 4 |
| G_and_S_frontomargin | R | 1 | 1 | 1 | 4 |
| G_and_S_occipital_inf | R | 2 | 4 | 3 | 4 |
| G_and_S_paracentral | R | 3 | 4 | 5 | 2 |
| G_and_S_subcentral | R | 2 | 1 | 5 | 5 |
| G_and_S_transv_frontopol | R | 2 | 1 | 2 | 2 |
| G_and_S_cingul-Ant | R | 3 | 1 | 1 | 1 |
| G_and_S_cingul-Mid-Ant | R | 2 | 1 | 3 | 1 |
| G_and_S_cingul-Mid-Post | R | 2 | 4 | 1 | 4 |
| G_cingul-Post-dorsal | R | 4 | 1 | 1 | 2 |
| G_cingul-Post-ventral | R | 4 | 4 | 4 | 1 |
| G_cuneus | R | 2 | 1 | 2 | 2 |
| G_front_inf-Opercular | R | 2 | 4 | 4 | 4 |
| G_front_inf-Orbital | R | 2 | 4 | 3 | 2 |
| G_front_inf-Triangul | R | 2 | 3 | 5 | 2 |
| G_front_middle | R | 2 | 4 | 4 | 4 |
| G_front_sup | R | 4 | 3 | 4 | 2 |
| G_Ins_lg_and_S_cent_ins | R | 2 | 1 | 4 | 1 |
| G_insular_short | R | 4 | 1 | 1 | 4 |
| G_occipital_middle | R | 4 | 4 | 4 | 4 |
| G_occipital_sup | R | 4 | 4 | 4 | 4 |
| G_oc-temp_lat-fusifor | R | 4 | 4 | 4 | 1 |
| G_oc-temp_med-Lingual | R | 1 | 4 | 2 | 3 |
| G_oc-temp_med-Parahip | R | 5 | 4 | 4 | 5 |
| G_orbital | R | 5 | 5 | 1 | 5 |
| G_pariet_inf-Angular | R | 2 | 1 | 1 | 2 |
| G_pariet_inf-Supramar | R | 2 | 3 | 1 | 2 |
| G_parietal_sup | R | 2 | 1 | 1 | 2 |
| G_postcentral | R | 3 | 4 | 4 | 4 |
| G_precentral | R | 4 | 4 | 4 | 4 |
| G_precuneus | R | 2 | 3 | 1 | 2 |
| G_rectus | R | 5 | 4 | 4 | 2 |
| G_subcallosal | R | 4 | 4 | 4 | 2 |
| G_temp_sup-G_T_transv | R | 3 | 1 | 4 | 3 |
| G_temp_sup-Lateral | R | 1 | 4 | 4 | 4 |
| G_temp_sup-Plan_polar | R | 4 | 1 | 4 | 4 |
| G_temp_sup-Plan_tempo | R | 3 | 3 | 4 | 4 |
| G_temporal_inf | R | 2 | 1 | 5 | 4 |
| G_temporal_middle | R | 4 | 4 | 5 | 2 |
| Lat_Fis-ant-Horizont | R | 3 | 3 | 3 | 3 |
| Lat_Fis-ant-Vertical | R | 3 | 3 | 3 | 4 |
| Lat_Fis-post | R | 1 | 3 | 2 | 1 |
| Pole_occipital | R | 1 | 1 | 2 | 2 |
| Pole_temporal | R | 1 | 3 | 2 | 2 |
| S_calcarine | R | 1 | 4 | 4 | 1 |
| S_central | R | 1 | 4 | 1 | 1 |
| S_cingul-Marginalis | R | 1 | 1 | 5 | 1 |
| S_circular_insula_ant | R | 3 | 1 | 2 | 1 |
| S_circular_insula_inf | R | 1 | 1 | 4 | 3 |
| S_circular_insula_sup | R | 1 | 1 | 1 | 1 |
| S_collat_transv_ant | R | 1 | 1 | 4 | 1 |
| S_collat_transv_post | R | 3 | 1 | 1 | 4 |
| S_front_inf | R | 3 | 5 | 5 | 5 |
| S_front_middle | R | 3 | 3 | 2 | 3 |
| S_front_sup | R | 3 | 1 | 4 | 4 |
| S_interm_prim-Jensen | R | 5 | 1 | 2 | 5 |
| S_intrapariet_and_P_trans | R | 3 | 3 | 1 | 3 |
| S_oc_middle_and_Lunatus | R | 5 | 3 | 5 | 3 |
| S_oc_sup_and_transversal | R | 1 | 4 | 2 | 3 |
| S_occipital_ant | R | 3 | 3 | 3 | 3 |
| S_oc-temp_lat | R | 1 | 1 | 1 | 3 |
| S_oc-temp_med_and_Lingual | R | 1 | 3 | 4 | 1 |
| S_orbital_lateral | R | 5 | 3 | 1 | 4 |
| S_orbital_med-olfact | R | 1 | 1 | 3 | 3 |
| S_orbital-H_Shaped | R | 1 | 1 | 2 | 3 |
| S_parieto_occipital | R | 1 | 4 | 1 | 3 |
| S_postcentral | R | 1 | 1 | 2 | 2 |
| S_pericallosal | R | 3 | 3 | 1 | 3 |
| S_precentral-inf-part | R | 3 | 3 | 1 | 3 |
| S_precentral-sup-part | R | 2 | 4 | 1 | 1 |
| S_suborbital | R | 1 | 3 | 2 | 4 |
| S_subparietal | R | 1 | 3 | 2 | 1 |
| S_temporal_inf | R | 3 | 4 | 5 | 5 |
| S_temporal_sup | R | 3 | 3 | 3 | 1 |
| S_temporal_transverse | R | 3 | 1 | 3 | 4 |

CTNs, cortical thickness networks; FDNs, fractal dimension networks; GINs, gyrification index networks; SDNs, sulcal depth networks.
