## Supplementary material for "Towards understanding interindividual differences in cortical morphological brain networks": Table S4

**Table S4**. Between-layers overlapping (dice coefficients) in community or module composition

|  | Module 1 | Module 2 | Module 3 | Module 4 | Module 5 |
| --- | --- | --- | --- | --- | --- |
| CTNs-FDNs | 0.416 | 0.063 | 0.469 | 0.333 | 0.375 |
| CTNs-GINs | 0.337 | 0.109 | 0.387 | 0.540 | 0.296 |
| CTNs-SDNs | 0.462 | 0.448 | 0.382 | 0.305 | 0.417 |
| FDNs-GINs | 0.380 | 0.080 | 0.339 | 0.529 | 0.316 |
| FDNs-SDNs | 0.337 | 0.054 | 0.366 | 0.386 | 0.500 |
| GINs-SDNs | 0.312 | 0.300 | 0.192 | 0.400 | 0.370 |
